# Reduction of the HIV-1 reservoir in T cells from persons with HIV-1 on suppressive antiretroviral therapy using expanded natural killer cells *ex vivo*

**DOI:** 10.1101/2022.09.05.506592

**Authors:** Mary Ann Checkley, Benjamin G. Luttge, Curtis Dobrowolski, David N. Wald, Deborah McMahon, Ghady Haidar, Michele D. Sobolewski, P. Nathan Enick, Joshua Cyktor, John W. Mellors, Jonathan Karn

**Affiliations:** Department of Molecular Biology and Microbiology, Case Western Reserve University School of Medicine, Cleveland, OH 44106, USA; Department of Pathology, Case Western Reserve University School of Medicine, Cleveland, OH 44106, USA; Department of Medicine, Division of Infectious Diseases, University of Pittsburgh School of Medicine, Pittsburgh, PA 15213, USA; Louis Stokes Cleveland VA Medical Center, Department of Pathology, Cleveland, OH 44106, USA

## Abstract

Treatment with latency-reversing agents (LRAs) alone has been ineffective in reducing HIV-1 reservoirs in persons with HIV-1 (PWH) on antiretroviral therapy (ART), due to inefficiencies in reservoir reactivation and adaptive immune responses. However, NK cells that are activated with cytokines may be able to target HIV-1 reservoirs more efficiently. To study the therapeutic potential of NK cells, we expanded blood NK cells from multiple donors *ex vivo* into CD56^bright^CD16^+^ “eNK” cells using artificial antigen presenting cells (aAPCs) expressing membrane-bound IL21. eNK cells express multiple activating receptors and are highly cytotoxic against specific target cells. eNK cells can also kill HIV-infected CD4 T cells via antibody dependent cell-mediated cytotoxicity (ADCC) using broadly neutralizing antibodies against HIV-1 Env gp120/gp41. Importantly, eNK cells from PWH on ART efficiently killed autologous HIV-1^+^ T cells reactivated by a combination of vorinostat (SAHA) and IL-15 or an IL-15 superagonist (N-803), as detected by declines in proviral load, inducible HIV-1 mRNA, and virus release. Adoptive immunotherapy with eNK cells is therefore a promising approach to reduce the latent HIV-1 reservoir in PWH when combined with LRA treatment.

**Author Summary:** Successful antiretroviral therapy (ART) eliminates progression to AIDS by reducing HIV to nearly indetectable levels by routine clinical measurements of blood samples. However, more sensitive DNA and RNA measurements show that most persons on ART retain a reservoir of long lived latently infected cells, which remain undetected by the immune system while no HIV is being produced. In nearly all cases, ART interruption results in a rebound of HIV production and spread, requiring an immediate return to ART. Currently the goal of HIV eradication is to achieve a “functional cure”, where HIV reservoirs are reduced to the point where ART can be interrupted indefinitely, and low levels of infected cells remaining can be controlled by the immune system. Our eradication strategy combines HIV latency-reversing agents (LRAs), ex vivo expansion of natural killer (NK) cells, and enhancement of specificity and killing of infected cells with broadly neutralizing antibodies against HIV. In this study, we have demonstrated that NK cells from person living with HIV can be isolated and expanded ex vivo into “eNK” cells that kill HIV-infected cells without killing uninfected cells, especially when broadly neutralizing antibodies are present, and can significantly reduce HIV reservoirs after LRA treatment.

## Introduction

Antiretroviral therapy (ART) is highly effective at suppressing the replication of infectious HIV-1 but does not diminish the reservoirs of latently infected cells, which persist throughout the body by evading immune detection. These reservoirs comprise mainly of transcriptionally silent, long-lived, memory CD4^+^ T cells, which are largely invisible to the immune system due to the lack of HIV-1 antigens [1]. In “Shock or Kick and Kill” strategies for HIV eradication, many have proposed that induction of HIV transcription with small molecule latency-reversing agents (LRAs) will help target latently infected cells for elimination by the immune system [2, 3]. Treatments with LRAs alone have failed to reduce HIV reservoirs in clinical trials [4–13]. This lack of viral clearance may be partly due to the accumulation of cytotoxic T lymphocyte (CTL) escape mutants in latent viral reservoirs [14]. Furthermore, CTLs from persons with HIV-1 (PWH) on ART are defective in killing autologous HIV-1+ resting primary CD4 T cells *in vitro* [15]. Thus, a successful HIV eradication strategy will likely require enhancement of innate immunity to efficiently target latent reservoirs after proviral reactivation with LRAs.

Activated natural killer (NK) cells may have the potential to meet the challenge of HIV-1 eradication. NK cells (defined as being CD3^-^CD56^+^) constitute 2-18% of lymphocytes in peripheral blood and generally fall into two main subsets: ∼90% CD56^dim^ and ∼10% CD56^bright^ [16]. The main effector function of the CD56^dim^ subset is induction of apoptosis in target cells, via release of cytotoxic granules or expression of death receptor ligands [17]. CD56^dim^ NK cells also typically express an Fcγ III receptor (CD16) that mediates antibody-dependent cell-mediated cytotoxicity (ADCC). This is a potentially important mechanism for killing of HIV-infected cells during acute infection. [18–21] It has been shown that broadly neutralizing antibodies (bNAbs), which are made in HIV-1 infected individuals and can neutralize the majority of HIV-1 isolates, can elicit ADCC [22]. Furthermore, infusion of bNAbs suppresses viremia, blocks infection and delays viral rebound in both non-human primates and in a humanized mouse model, and bNAbs have also been shown to delay viral rebound after ART interruption in persons living with HIV [23–34]. In contrast, the main effector function of CD56^bright^ NK cells is to secrete cytokines and chemokines that modulate the immune response rather than to directly elicit cytotoxicity. However, even these immunoregulatory NK cells can become cytotoxic after activation with cytokines [35–38]. As part of the innate immune system, NK cells sense potential target cells by expressing an array of activating and inhibitory receptors, rather than by antigen recognition. This also helps NK cells kill target cells that otherwise would escape CTL recognition [39]. Cytolytic activity is only activated if the balance of ligated activating receptors on NK cells outweighs the binding of inhibitory receptor ligands that are normally expressed on healthy cells. By this mechanism, expression of some HIV-1 proteins is sufficient to trigger recognition by NK cells. For example, HIV-1 Nef and Vpu downregulate MHC-I in infected cells. This results in some immune suppression by inhibiting antigen presentation but also activates NK cells by decreasing the ligation of MHC-I to NK cell-inhibitory receptors [40–43]. Additionally, NK cells have activating receptors that specifically recognize ligands upregulated by HIV Vpr and other signs of stress [44].

In this study, we investigated whether we could expand NK cells from PWH ex vivo to enhance their cytotoxicity against HIV-infected cells and significantly deplete HIV reservoirs in vitro. The clinical applications of this approach are that autologous, expanded NK cells could serve as effectors against the HIV reservoir in vivo by adoptive transfer, using methods originally developed as immunotherapy for cancer. One potential concern is that there is an expansion of dysfunctional CD56^-^ NK cells in chronic HIV infection, which diminishes the availability of circulating cytotoxic NK cells [45–47]. However, even NK cells isolated from peripheral blood of HIV-negative donors have relatively low cytotoxicity in the absence of activation [48]. This lack of function can be overcome *in vitro* by treatment with cytokines such as IL-2, IL-15, or IL-15 derivatives (N-803) [49–53]. Another potential challenge is that cytotoxicity can still be limited if there is an insufficient effector to target cell ratio, and NK cells isolated from peripheral blood are not abundant enough to provide a therapeutic dose, even in HIV-negative donors. Fortunately, the development of artificial antigen presenting cell (aAPC) technology now allows massive expansion of cytotoxic “eNK” cells from a simple blood draw [54]. Here we utilized aAPCs expressing membrane-bound IL-21 (mbIL21) to expand eNK cells from peripheral blood mononuclear cells (PBMCs) and systematically analyzed their expression of inhibitory and activating receptors, cytotoxicity, and ADCC using bNAbs against autologous HIV-infected cells [48, 55]. Using leukapheresis products from PWH on ART, we found that autologous eNK cells can significantly reduce the latent reservoir *in vitro* after proviral reactivation, as measured by losses of proviral load, virus release, and inducible cell-associated HIV mRNA. We also expanded autologous eNK cells using Good Manufacturing Practice (GMP) protocols with an FDA-approved aAPC line (NKF), which yielded similar results [56]. Our results suggest that translation of this approach to adoptive transfer of eNK cells for targeting HIV reservoirs *in vivo* could become an important component of an HIV cure strategy.

## Results

### Expansion of NK cells from HIV^+^ participants using feeder cells expressing mbIL21 yields a highly activated phenotype

Given that chronic HIV infection has been shown to reduce populations of more cytotoxic NK cell subsets (CD56^dim^ CD16^+^) [57], we explored whether it would be feasible to expand and activate autologous cytotoxic NK cells from PWH *ex vivo* to enhance NK cell function by adoptive immunotherapy. Using frozen leukopaks, we enriched NK cells from well-suppressed ART-treated PWH and from persons without HIV-1. Using aAPC technology, each NK cell isolate was cocultured with K562 feeder cells expressing membrane-bound IL21 (C9.K562-mbIL21) for approximately 3-4 weeks, resulting in a 150- to 500-fold expansion of eNK cells. NK cell isolates were analyzed by flow cytometry to evaluate the expression of CD56, CD16, and other NK cell receptors before and after expansion (Figs 1A-B). Prior to expansion, the phenotype of isolated NK cells was primarily CD56^dim^CD16^+^ or CD56^-^CD16^-^, with the CD56^-^ subset being more prevalent in PWH. By contrast, the expanded NK (eNK) cells were predominantly CD56^bright^ (76-98%), regardless of HIV status.

**Fig 1.**
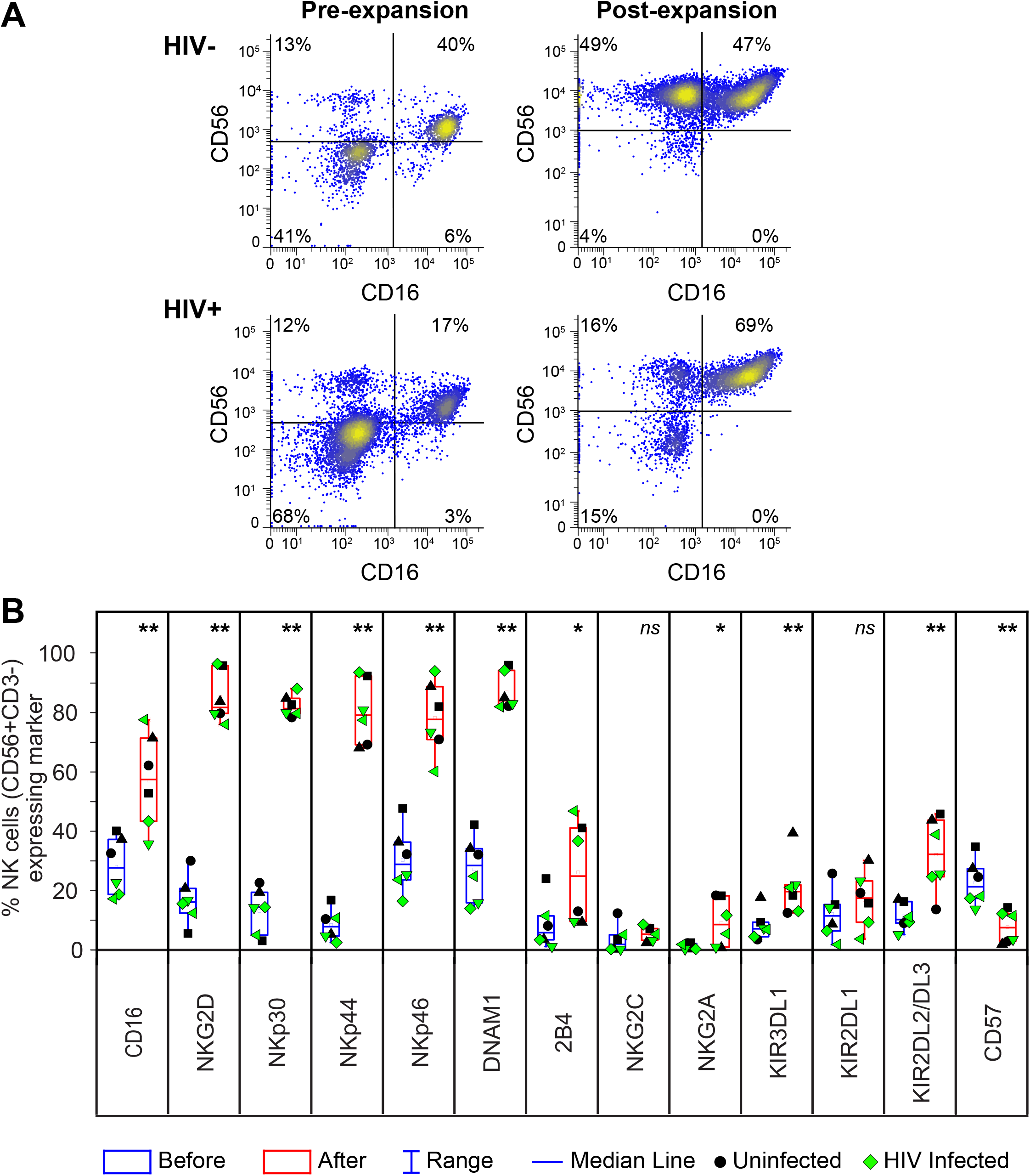
Expansion of NK cells from PWH yields NK cells that express high levels of activating receptors. **(A-B)** NK cell phenotypic analysis by flow cytometry before and after expansion with irradiated C9.K562.mbIL21 cells. **(A)** Representative dot plots gated on CD3-cells showing expression of NK cell markers CD56 and CD16 before and after (pre- and post-) expansion from representative HIV- and HIV+ donors. **(B)** Frequency of CD56+ NK cells expressing NK cell markers before (blue box) and after expansion (red box) from HIV- (black symbols) and HIV+ (green symbols) donors (*n* = 3 biological replicates per group). Significant differences after expansion were determined using paired *t*-tests, \**p*<0.05 2B4, \*\**p*<0.01, and ns, non-significant.

The proportions of CD56^+^ eNK cells expressing key NK cell-activating receptors were significantly higher after expansion, including CD16 (2-fold), NKG2D (7-fold), NKp30 (11-fold), NKp44 (15-fold), NKp46 (3-fold), DNAM1 (4-fold), and 2B4 (5-fold) (Fig 1B). This directly correlated with higher expression levels per cell for each of these receptors, as shown by mean fluorescence intensity (MFI) (S1A Fig). The proportion of CD56^+^ eNK cells expressing inhibitory receptors was also modestly increased, including KIR3DL1 (3-fold), KIR2DL2/DL3 (3-fold), and NKG2A (8-fold). Interestingly, eNK cells were predominantly negative for CD57, which is a marker of NK cell maturation associated with decreased proliferative capacity and responsiveness to cytokines [58]. To further evaluate the potential of this protocol for immunotherapy by adoptive transfer of NK cells, we also examined the expansion of NK cells with an alternative aAPC cell line expressing mbIL-21 (NKF cells) that has been certified for clinical use as an investigational new drug (IND) [56]. This yielded eNK cells with a very similar phenotype to eNK cells expanded with C9.K562-mbIL21 (S1B Fig). In summary, expansion of NK cells from HIV^+^ participants using aAPCs expressing mbIL21 consistently yields predominantly CD56^bright^ eNK cells that express high levels of activating receptors necessary for cytotoxicity while maintaining several inhibitory receptors that are important for regulation.

### eNK cells are highly cytotoxic and compatible with LRAs

We then determined whether NK cells expanded from PWH donors are cytotoxic, relative to eNK cells from HIV negative donors, using K562 target cells. We first demonstrated cytotoxicity using a PanToxiLux substrate (PS) that detects NK cell-mediated delivery of granzyme B into target cells (Fig 2A and S2A-B Fig). We also developed a K562-RFP cell line that can detect cytotoxicity by loss of RFP, which we show directly correlates with cell death, based on fixable viability dye staining (FVD) (Fig 2B and S2C Fig). Using a 2:1 (eNK:K562) ratio, eNK cells targeted an average of 55-59% of K562 cells for killing detected by PS cleavage. In average about 40-45% of eNK cell input from HIV- and HIV+ donors was required to kill 50% of K562 target cells as detected using the K562-RFP killing assay. By either approach, eNK cells from PWH demonstrated a highly significant degree of cytotoxicity, which was equivalent to that of eNK cells from HIV-negative donors. These results suggests that NK cells from PWH can be efficiently expanded into highly cytotoxic eNK cells using aAPCs.

**Fig 2.**
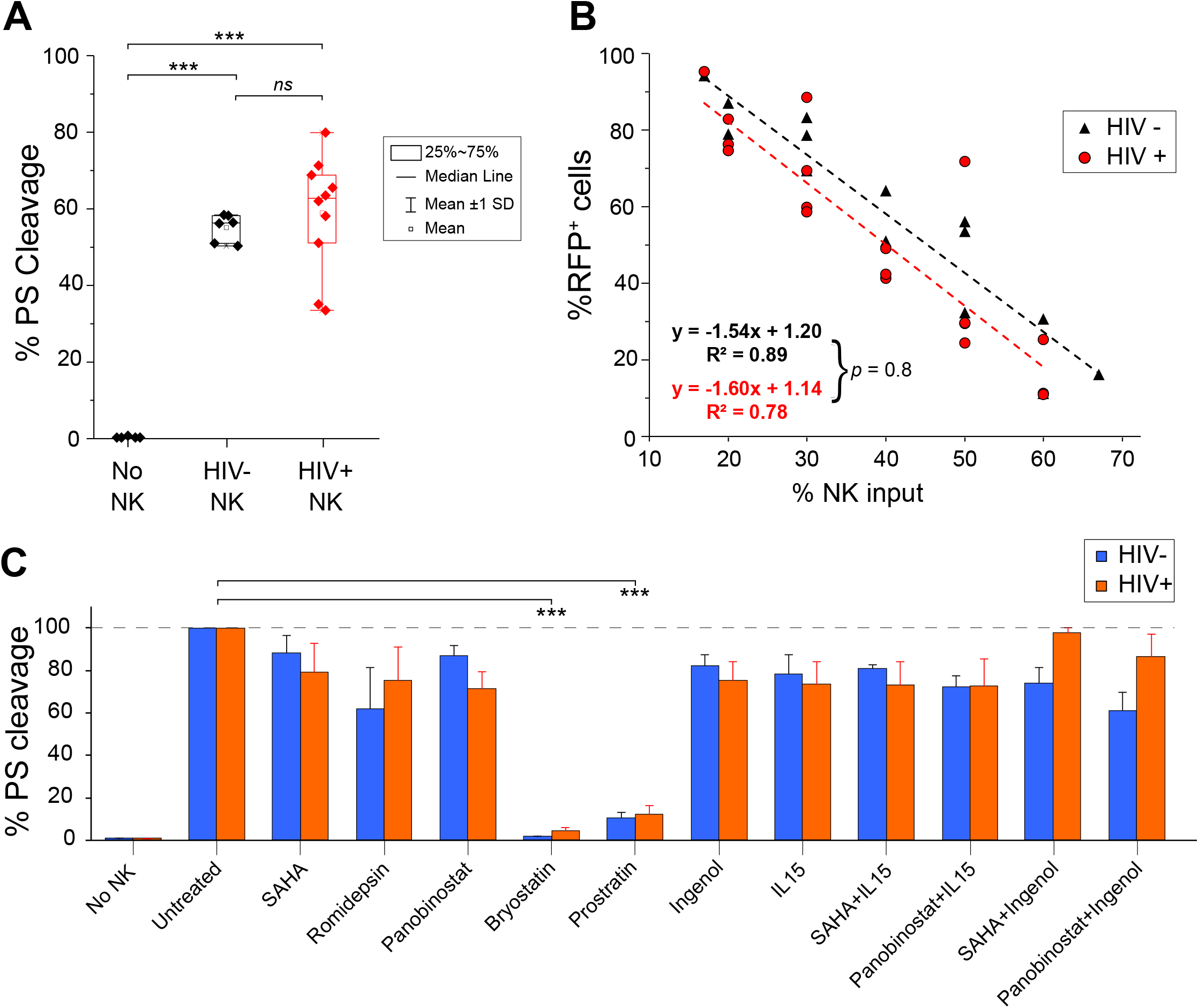
eNK cells from PWH are highly cytotoxic against K562 target cells and LRA effect on their cytotoxicity. **(A-C)** eNK cells from HIV+/-donors were cocultured with K562 cells and killing was measured by flow cytometry. **(A)** Killing of K562 target cells by eNK cells at a 2:1 (E:T) ratio is measured by PanToxiLux Substrate (PS) cleavage. The percentage of K562 cells with cleaved PS is shown. *n* = 5 biological replicates for HIV-negative group and *n = 10* biological replicates for HIV-positive group. Each sample was performed in triplicate. **(B)** CTV-stained eNK cells were cocultured with K562-RFP target cells. Cytotoxicity was then detected in target cells (CTV-events) by loss of RFP. *n* = 3 biological replicates per group. Each sample was performed in triplicate. There was no significant difference in cytotoxicity between eNK cells from donors with or without HIV (*p* = 0.8). **(C)** LRA effect on eNK cell cytotoxicity. eNK cells from HIV+/-donors were treated with LRAs for 24 h or left untreated, then washed and cocultured with K562 cells at a 2:1 (E:T) ratio to measure cytotoxicity by PS cleavage. LRAs were used alone or in combination as follows: SAHA (335 nM), romidepsin (10 nM), panobinostat (20 nM), bryostatin (5 nM), prostratin (1 µM), ingenol (100 nM), and IL15 (10 ng/ml). Percentage of K562 cells targeted for killing (% PS cleavage) by eNK cells relative to no LRA treatment (untreated) is shown. *n* = 2 biological replicates from 3 independent experiments for HIV-negative group and *n* = 3 biological replicates for HIV-positive group. Error bars show SEM. Significant differences were determined using two-tailed unpaired *t*-tests of the means in panel A, or the slopes created by linear regression in panel B, and paired *t*-tests in panel C, *** *p*<0.001.

To visualize eNK cell cytotoxicity in the presence of HIV protein expression, we analyzed 3C9 cells (Jurkat cells transduced with a latent HIV-GFP construct) by time lapse microscopy. TNFα-activated 3C9 cells were mixed with a dead-cell stain (Sytox Orange) and imaged at 37 °C, either alone or with fluorescent eNK cells. In the absence of eNK cells, 3C9 cells remained evenly distributed and no cell death was detectable (S1 Movie). In contrast, eNK cells were highly motile and conjugated target cells within 10 minutes (S2D Fig). Using a 1:4 (effector:target cell) ratio, approximately 6-12% of target 3C9 cells were typically killed within 45 minutes (S2D Fig and S2 Movie). Some individual eNK cells killed up to four target cells during this time, which demonstrates their capacity for serial killing. Our results show that eNK cells are highly motile killers that rapidly conjugate multiple target cells and elicit cytotoxicity without self-injury.

Given that NK cells would be exposed to LRAs during a cure treatment using the shock and kill approach, we also evaluated the effect of clinically relevant concentrations of histone deacetylase inhibitors (HDACi, like SAHA, panobinostat, romidepsin) and protein kinase C (PKC) agonists (bryostatin, prostratin, and ingenol) on eNK cell-mediated cytotoxicity (Fig 2C). Since the cytokine IL-15 activates both NK and T cells, and can enhance the reversal of latency by HDACi, we also tested the cytotoxicity of eNK cells after treatment with IL-15 alone or in combination with SAHA or panobinostat. Treatment of eNK cells overnight with any HDAC or with ingenol led to only a modest decrease in cytotoxicity against K562 cells, which was not significantly affected by the addition of IL-15. The combination of ingenol with HDACi was most compatible with eNK cells from HIV+ donors. In contrast, overnight treatment of eNK cells with bryostatin or prostratin dramatically reduced their cytotoxicity. Thus, any HIV reactivation strategy using LRAs should also take into account the potential impact on NK cell effector functions.

### eNK cells specifically kill acutely HIV-1 infected autologous T cells

To evaluate whether eNK cells would specifically kill HIV-1-infected T cells in a pool of uninfected cells, we used autologous CD4^+^ memory T (T_M_) cells at 3-4 days after infection with R5-tropic, replication-competent HIV-GFP. After treatment with IL-15 overnight, infection of T_m_ cells with HIV-GFP yielded 1-5% GFP^+^ cells (Fig 3A). For discrimination from target cells, eNK cells were stained with a concentration of CellTrace Violet (CTV) that does not affect NK cell cytotoxicity or viability. Killing of HIV^+^GFP^+^ T_m_ cells by coculture with CTV-eNK cells for 24 h was detected by loss of GFP, while general cytotoxicity was detected by staining with propidium iodide (PI). Under these conditions, we consistently observed that eNK cells killed 38% of HIV^+^GFP^+^ T_m_ cells on average when cocultured at an effector to target ratio (E:T) of 1:1 (50% NK cell input), and we observed 15% killing when using at an E:T ratio of only 1:9 (10% NK cell input) (Fig 3B). In contrast, off-target effects detected by PI staining of GFP^-^ cells were minimal (S3A Fig), which accounts for the highly specific killing of HIV^+^ cells.

**Fig 3.**
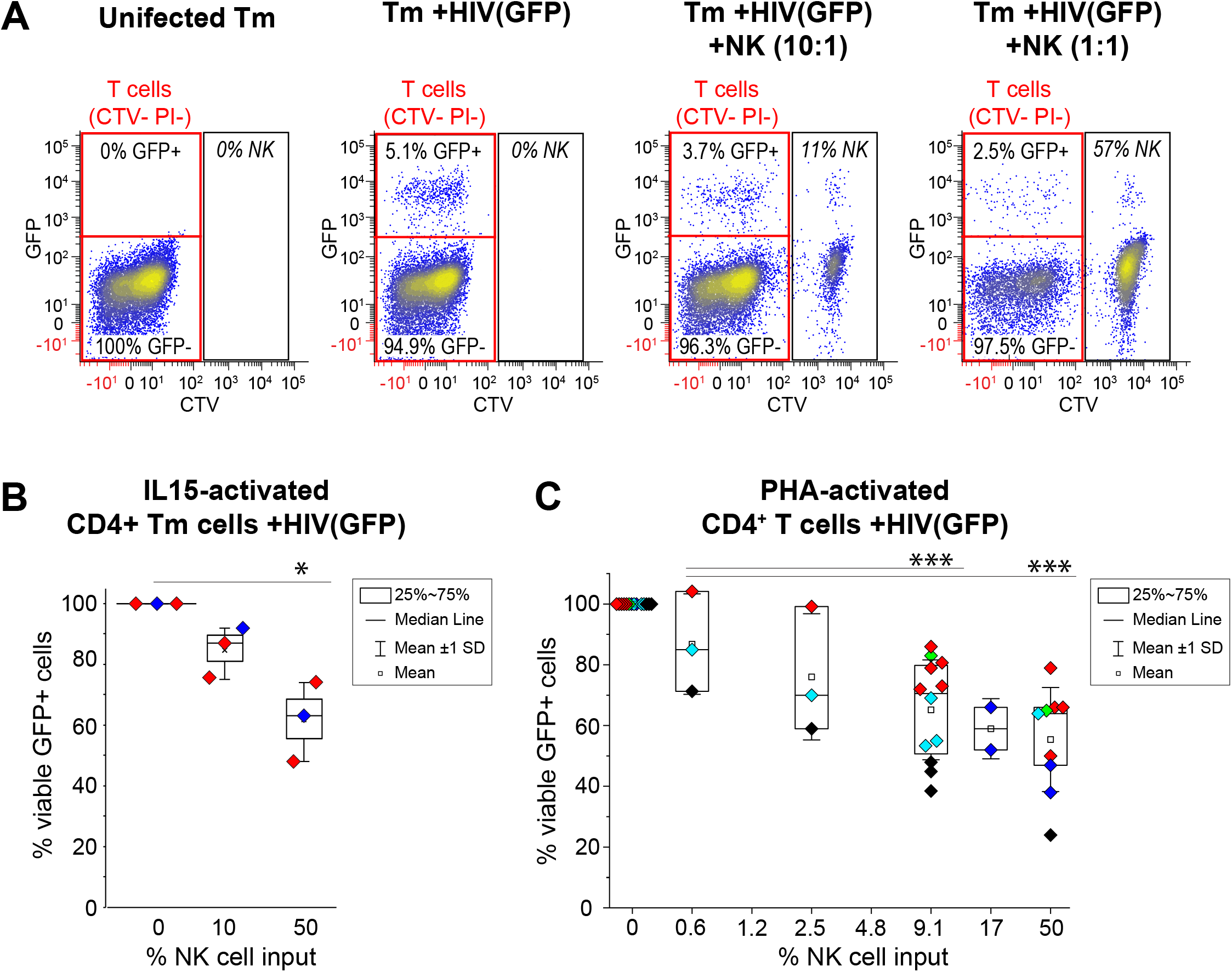
eNK cells specifically and efficiently kill acutely HIV-1-infected autologous T cells. **(A-C)** CTV-stained eNK cells were cocultured with autologous primary T cells infected with replication-competent HIV expressing GFP [HIV(GFP)]. Using flow cytometry, killing of HIV-infected T cells was measured by loss of GFP and total target cell death was detected by PI staining of CTV-cells. **(A)** Representative dot plots of uninfected memory T cells (T_m_), HIV(GFP)-infected T_m_ cells (T_m_ +HIV(GFP)) alone or after 1 day coculture with eNK cells at a 1:10 and 1:1 (NK:T_m_) ratio. Percentages of GFP+ and GFP-T_m_ cells are highlighted by red gates. CTV+ eNK cells are shown, but excluded from analysis. **(B-C)** Box plots showing dose-dependent, eNK cell-mediated killing of activated CD4 T cells infected with HIV(GFP). The percentage of viable GFP^+^ T_m_ cells (CTV^-^ PI^-^) in cultures with eNK cells was normalized to that of cultures without eNK cells. **(B)** eNK cell-mediated killing of IL15-activated memory T cells (T_m_) infected with HIV(GFP). *n* = 3 independent experiments, and each biological replicate (shown in unique colors) was performed in triplicate. **(C)** eNK cell-mediated killing of PHA-activated CD4+ T cells infected with HIV. *n* = 2-12 independent experiments for each dose of NK cells (symbols show mean of each sample performed in triplicate), using 5 biological replicates (unique color for each). Significant differences were determined using paired *t*-tests, \**p*<0.05, \*\*\**p*<0.001.

To compare IL15-activated cells with maximally activated target cells, PHA-activated CD4^+^ T cells were infected with HIV-GFP, which yielded 5-15% GFP^+^ cells within 3-4 days. These target cells were then cocultured with autologous eNK cells for 24 h. Similar to our results when using IL-15, we observed that autologous eNK cells kill 45% of PHA-activated HIV^+^GFP^+^ CD4 T cells on average with an E:T ratio of 1:1 (50% NK cell input) and 30% killing when using at an E:T ratio of 1:10 (9.1% NK cell input) (Fig 3C).

The specificity of eNK cell-mediated killing was also demonstrated using a modified PanToxiLux assay. With this approach, specific eNK cell-mediated killing of HIV^+^GFP^+^ cells correlated with loss of GFP, and overall cytotoxicity was detected by cleavage of a dual caspase-8/granzyme B substrate (PS) in target cells (S3B Fig). PHA-activated CD4 T cells were either acutely infected with replication-competent CXCR4-tropic or CCR5-tropic HIV-GFP [HIV(X4) and HIV(R5), respectively], or transduced with an HIV-1 gag^-^pol^-^ construct (PHR’) that expresses mCD8α-GFP at the cell surface (S3C Fig). Infection or transduction of PHA-activated CD4 T cells yielded 5-7% GFP^+^ T cells, which were then cocultured with autologous eNK cells. We observed significant NK cell-mediated killing of GFP+ CD4 T cells (∼42% of PHR’-transduced cells and ∼54% of HIV-GFP-infected cells) (S3D Fig). In all cases, there was <1% killing of uninfected (GFP^-^) cells, detected by PS cleavage (S3C Fig). Furthermore, expansion of NK cells with NKF cells also yielded highly cytotoxic eNK cells that specifically killed PHA-activated, autologous HIV(R5)-GFP-infected CD4 T cells (S3E Fig). Overall, these data suggest that eNK cells can specifically target and kill HIV^+^ cells with no collateral damage to uninfected cells.

### eNK cell-mediated killing of HIV-infected cells can be enhanced with ADCC

Given that expansion of NK cells with mbIL21 yields CD16+ eNK cells, we assessed whether eNK cells can kill HIV-infected cells via ADCC using a panel of broadly neutralizing antibodies (bNAbs) against HIV Env. To assess ADCC we chose bNAbs that target the V3 loop (PGT121, PGT126, PGT128, PGT135, 10-1074, and 2191), the CD4 binding site (VRC01, G54W, VRC03, CH106, 3BNC117, and VRC-CH31), the V1/V2 loops (PG9, PG16, PG145, HG107, and CH58), and gp41 or MPER (7B2, 7B2-AAA, 2F5, 7H6, and 10E8). PHA-activated CD4 T cells were acutely infected with replication-competent CXCR4-tropic or CCR5-tropic HIV-GFP (X4 and R5, respectively). Individual bNAbs were then added to target cells and cocultured with autologous eNK cells at an E:T ratio of 1:10. As negative controls for ADCC, eNK cells and target cells were cocultured with either a negative control antibody or with no antibody. bNAbs were considered capable of eliciting ADCC if they significantly enhanced NK cell-mediated killing relative to negative controls. Killing of HIV-GFP-infected cells was detected by loss of GFP. We found that bNAbs that target the CD4 binding site of HIV Env were able to elicit the ADCC function of eNK cells against cells infected with both HIV(R5) and (X4) (S4A Fig). As expected, bNAbs that target the N-glycan V3 loop were only able to elicit ADCC against cells infected with HIV(R5). Some of the antibodies that target N-glycan V1/V2 loops (PG16) and the MPER region (2F5 and 10E8) also elicited ADCC against both HIV(R5) and (X4)-infected cells. The negative control antibody, which is a human antibody that recognizes the H1N1 hemagglutinin protein of influenza A virus, does not elicit ADCC from eNK cells against CD4 T cells infected with HIV (S4B Fig). We then confirmed that ADCC could be consistently detected with eNK cells from multiple donors, using autologous CD4 T cells infected with HIV(R5) and selected bNAbs or with no antibody, and an E:T ratio of 1:10. As detected by loss of GFP+ target cells, NK cell-mediated killing was still evident even in the absence of antibody at this relatively low concentration of effector cells. Aside from PG16, each of the bNAbs tested showed a statistically significant enhancement of target cell killing (Fig 4). To test the potency of bNAbs eliciting ADCC, we titrated selected individual antibodies at a range of 0.16-10 ug/ml. We found that significant ADCC was detected over a wide range of antibody concentrations when using an E:T ratio of 1:10. In general, saturation of ADCC occurred between approximately 1-2.5 ug/ml. VRC01 was found to be the most potent, with a maximum of ADCC at 0.625 ug/ml (S4C Fig). Overall, we found that eNK cells have a significant capacity to kill HIV-infected CD4 T cells via ADCC when using bNAbs that target HIV Env.

**Fig 4.**
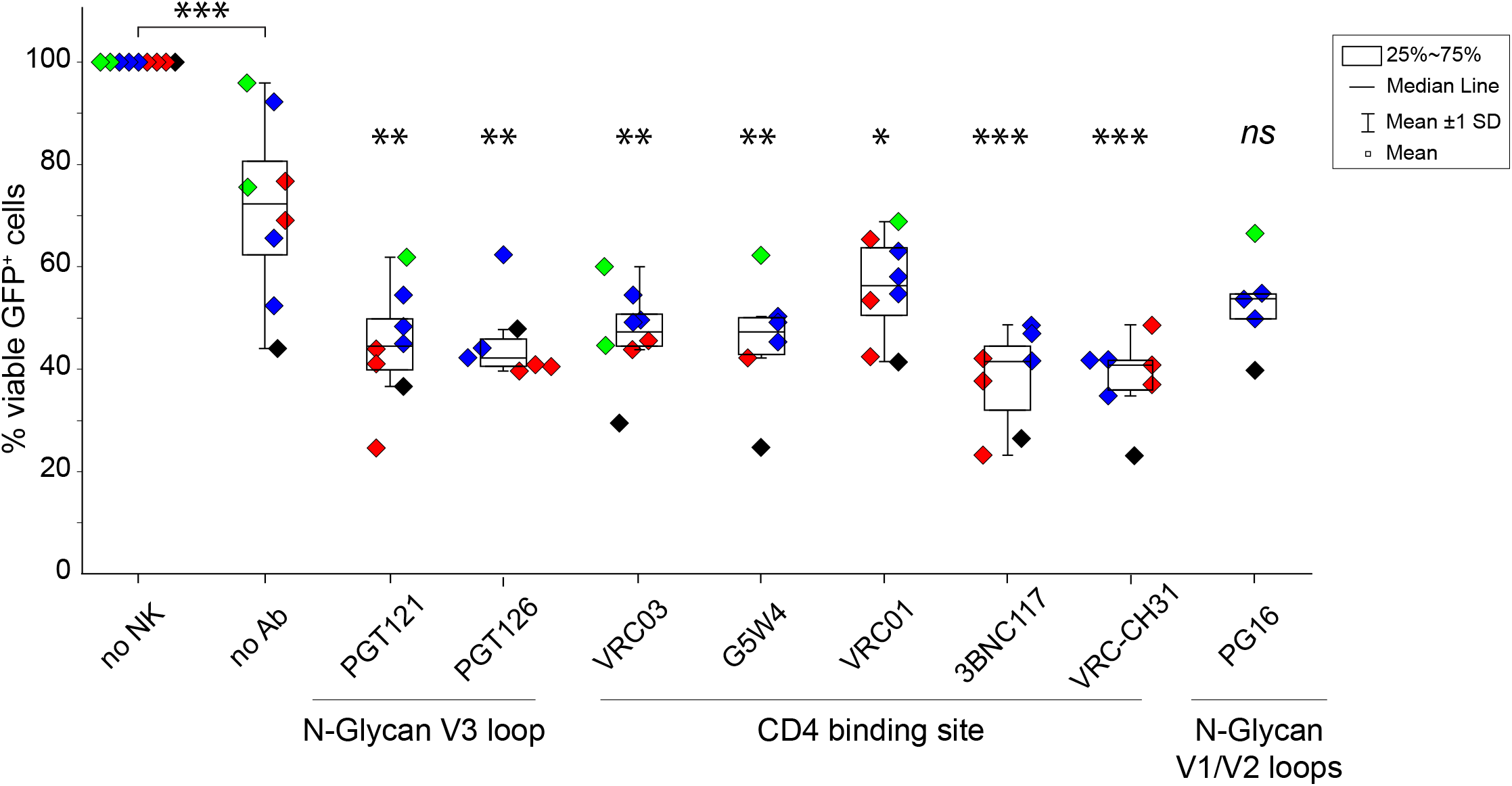
eNK cell-mediated killing of HIV-infected cells can be enhanced with ADCC using bNAbs against HIV Env. PHA-activated CD4+ T cells were acutely infected with replication-competent R5-tropic HIV(GFP) and cultured for 1 day with CTV-stained eNK cells at a 1:10 (NK:T) ratio with anti-HIV Env at 10 µg/ml and 10µM raltegravir. By flow cytometry, killing of HIV-infected T cells (CTV-) is measured by loss of GFP, and total target cell death is measured by PI staining. The percentage of viable GFP^+^ T cells (CTV^-^ PI^-^) in cultures with eNK cells was normalized to cultures without eNK cells. *n* = 4 biological replicates. Each donor is shown in a different color, and each symbol represents the mean of samples in triplicate from each independent experiment. Significant differences were determined using unpaired *t*-tests. \**p*<0.05,\*\**p*<0.01, \*\*\**p*<0.001. *ns*, non-significant.

### Autologous eNK cells from PWH reduce HIV release from CD4 T cells after latency reversal

We then wanted to determine whether autologous eNK cells can show antiviral activity against reservoirs of latently infected CD4 T cells from PWH after *in vitro* LRA treatment. As a preclinical model for HIV eradication, we combined Good Manufacturing Practice (GMP) for NK cell expansion, clinically used LRAs, and a clinical assay for HIV detection. NK cells were expanded from CD3-depleted PBMCs of PWH (Donors 1, 2, and 3) using an FDA-approved cell line (NKF) (Fig 5A). This resulted in ∼35-fold expansion of predominantly CD3^-^CD56^bright^CD16^+^ eNK cells with <0.1% T cell contamination after four treatments with irradiated NKF cells over a span of 4 weeks (S5A Fig). These GMP-eNK cells had a similar phenotype to eNK cells expanded with the C9.K562-mbIL21 aAPC (compare Fig 1A) and efficiently killed K562-RFP cells (S5B Fig). About 35-38.5% of GMP-eNK cell input from these donors was required to kill 50% of K562-RFP target cells which was comparable to NK cells expanded with C9.K562-mbIL21 aAPC (compare Fig 2B).

**Fig 5.**
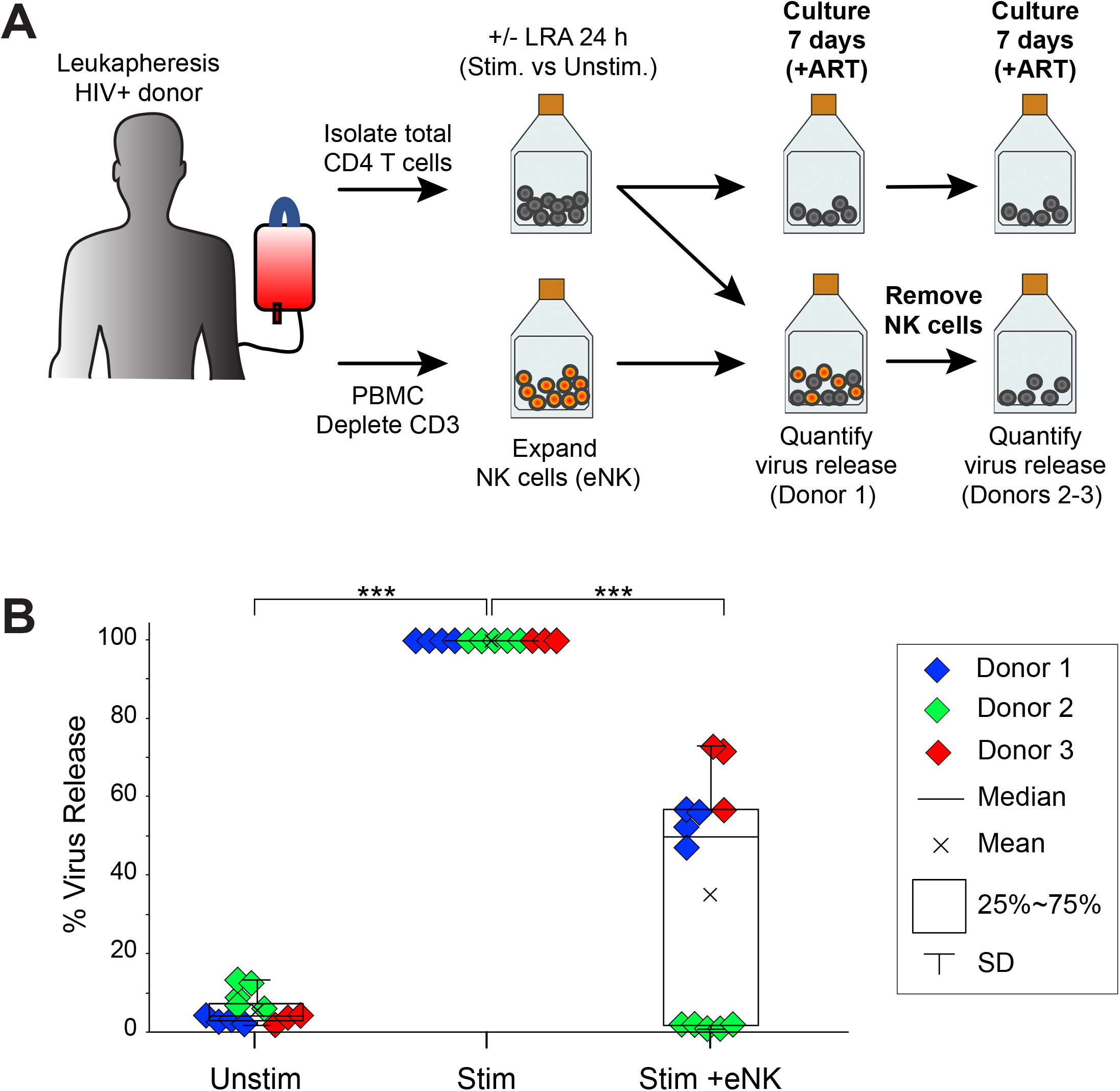
eNK cells from PWH reduce HIV release from autologous CD4 T cells after latency reversal. **(A)** Schema of eNK killing assays detected by virus release. CD4 T cells from ART-treated PWH were isolated from PBMCs and activated with LRA for 24 h. In parallel, NK cells were enriched from PBMCs by CD3 depletion and expanded into eNK cells using NKF cells. Cocultures of activated CD4 T cells were mixed with an equal number of autologous eNK cells and incubated for 7 days in growth media containing antiretroviral drugs. After the coculture, supernatants were collected to measure virus release (Donor 1). For Donors 2 and 3, eNK cells were removed at Day 7 and incubated for an additional 7 days prior to collection of supernatant. **(B)** Virus release measurements in CD4 T cells from PWH. Supernatants from CD4 T cells cultures alone or with eNK cells were collected to quantify HIV release by detecting HIV RNA in the supernatant using the Aptima HIV-1 Quant assay. After complete media change at Day 5 (Donor 1) and Day 8 (Donors 2-3), *de novo* virus release was detected at Day 7 (Donor 1) and Day 14 (Donors 2-3). Percent Virus Release was normalized to stimulated CD4 T cells with no NK cells (Stim) and compared to stimulated CD4 T cells with eNK cells (Stim+eNK). CD4 T cells with no stimulus (Unstim) were also tested as a control for HIV latency in PWH samples. *n* = 3 biological replicates. Significant differences were determined using paired *t*-tests. \*\*\**p*<0.001.

We then wanted to determine whether GMP-eNK cells can show antiviral activity against reservoirs of latently infected CD4 T cells from PWH by measuring HIV release after latency reversal (Fig 5A). Total CD4+ T cells were isolated from Donors 1-3, stimulated overnight to reactivate HIV via TCR stimulus or LRA treatment, and cocultured with an equal number of autologous eNK cells in the presence of antiretroviral drugs. As a positive control for reactivation, target cells from Donor 1 were treated with a TCR stimulus. For Donors 2 and 3, we utilized SAHA in combination with the IL-15 superagonist N-803, which has shown promising results *in vivo* in enhancing CD8+ T cell and NK cell function by targeting SIV reservoirs in B cell follicles and reversing HIV latency, which we have also demonstrated by detection of induced cell-associated HIV mRNA (S5C Fig) [59–62]. For all samples, release of HIV RNA was quantified using the Aptima HIV-1 Quant assay after coculture with eNK cells. For Donor 1, culture media was replaced on Day 5 and analyzed on Day 7. For Donors 2 and 3, eNK cells were removed on Day 7 by CD56^+^ depletion (S5D Fig), and the remaining T cells were cultured for an additional 7 days prior to analysis. Virus release was normalized to stimulated CD4 T cells with no NK cells (Stim) and compared to stimulated CD4 T cells with eNK (Stim+eNK) (Fig 5B). Samples with no stimulus (Unstim) were also were cultured in parallel as a control. HIV release from unstimulated cells was minimal for all donors, as expected from latent HIV reservoirs, and virus release after stimulation was highly significant. In contrast, stimulated CD4 T cells incubated with eNK cells consistently showed a decrease in virus release, which ranged from a modest 1.5 to 2-fold reduction (Donors 1 and 3) to nearly complete loss of virus release (Donor 2). Our results suggest a significant antiviral activity against reactivated HIV+ CD4 T cells from PWH when cultured with eNK cells.

### eNK cells from PWH reduce proviral loads in autologous CD4 T cells after latency reversal

We developed a novel proviral assay that includes the detection of potentially intact proviral load (PIPL) by duplex digital PCR (Fig 6). Specifically, by combining TaqMan probe/pair sets to detect both *gag* and *env* in the same sample, we can evaluate effects on single-positive *gag^+^*(*env*^-^) and *env^+^*(*gag*^-^) proviruses that are missing a portion of *env* or *gag*, rendering them potentially defective, as well as evaluating potentially intact, double-positive (*gag*^+^*env*^+^) proviruses. Using DNA from memory CD4 T cells, we define the number of *gag*^+^*env*^+^ proviruses per million cells as the potentially intact proviral load (PIPL) and the percent of *gag*^+^*env*^+^ proviruses relative to the total is the % PIPL. Unlike conventional proviral assays that rely on a single probe and primer pair to quantify proviral load, analysis of both defective and potentially intact proviruses provides a more complete picture of the HIV reservoir. A similar approach for an intact proviral DNA assay (IPDA) using duplex digital droplet PCR has been recently described [63, 64]. Due to the potential for PCR inhibition at high DNA concentrations, which might lead to a false detection of *gag*^-^ or *env*^-^ proviruses, we confirmed that detection of intact proviruses in our assay is not restricted by the excess of uninfected cellular DNA in primary samples. For this control, we tested pNL4-3 mixed with an excess of HIV-negative cellular DNA, which yielded exclusively *gag*^+^*env*^+^ signals (>99% PIPL). In comparison, the proviral load of CD4 memory T (T_m_) cell DNA from a well-suppressed ART-treated HIV^+^ donor comprised predominantly single-positive (defective) proviruses, and *gag*^+^*env*^+^ proviruses were rare (∼4% PIPL) (Fig 6A).

**Fig 6.**
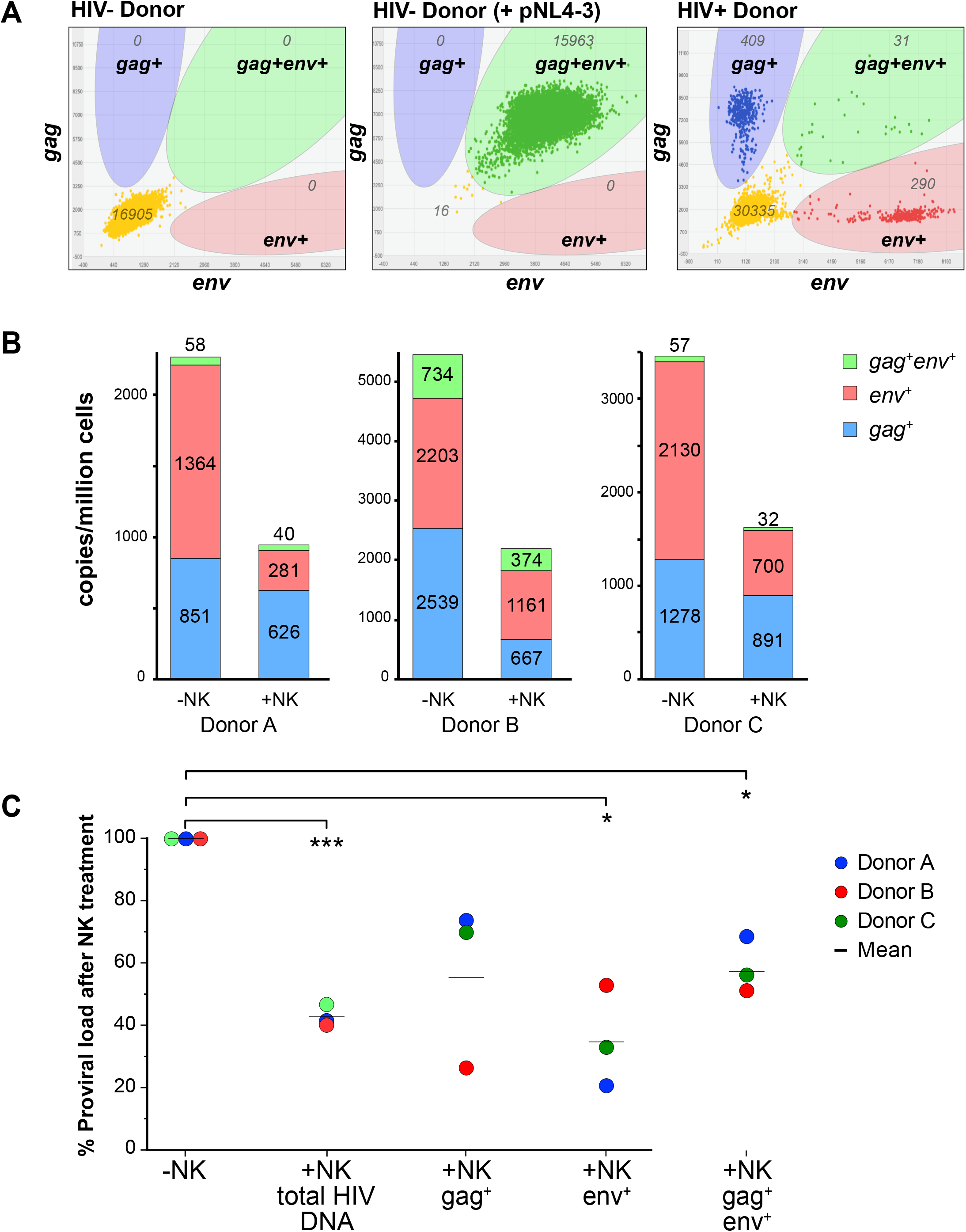
Loss of proviral load in memory T cells from ART-treated PWH after treatment with autologous eNK cells. **(A)** Quantification of HIV-1 proviral load by duplex digital PCR (PIPL assay). HIV-1 proviruses were counted by QuantStudio 3D digital PCR using FAM-labeled *gag* and VIC-labeled *env* probes in duplex. PBMC DNA from an HIV-negative donor (left) is a negative control, which when mixed with a trace amount of pNL4-3 is a control for fully intact proviruses (middle). Memory T cell DNA from an HIV+ donor on ART (right) contains an excess of *gag+* (*env*-negative) proviruses and *env+* (*gag*-negative) HIV proviruses, with a limited number of potentially intact (*gag^+^env^+^*) proviruses. **(B)** Indirect detection of NK cell-mediated cytotoxicity by quantifying proviral loads (HIV DNA copies per million cells). CD4 memory T cells (T_m_) isolated from PWH were treated overnight with SAHA+IL15 (LRA) prior to culture alone (-NK) or 1:1 with eNK cells (+NK) for 7 days. After reisolation of T_m_ cells by CD3+ selection, cellular DNA was analyzed by the PIPL assay. Contributions of *gag^+^*, *env^+^*, and *gag^+^env^+^* proviruses to the total is shown for each sample. **(C)** Statistical analysis of NK cell-mediated effects on proviral loads. Data from PIPL assays of NK cell-treated T_m_ cells (% proviral load after NK treatment) were normalized to T_m_ cells cultured alone (-NK) and averaged. Total HIV DNA is the sum of all proviruses detected with either probe. Relative changes in defective (*gag^+^ or env^+^*) or potentially intact (*gag^+^env^+^*) proviral loads are shown. *n* = 3 biological replicates. Statistical differences in the means of proviral loads +/-NK cells were determined by paired, two-tailed Student’s *t* tests (*two-tailed *P* value < 0.05, *** *P* < 0.001).

We then applied our PIPL assay to quantify changes in proviral load after treatment with NK cells. After 7 days of culturing LRA-treated T_m_ cells with eNK cells, T_m_ cells were reisolated by positive selection against NK cells (S5A Fig) and cellular DNA was analyzed. On average, there was a highly significant loss of total proviral load after incubation with eNK cells (57% reduction, *p*<0.001), including a significant loss in *env*^+^ cells (65% reduction, *p*<0.05) and *gag*^+^*env*^+^ cells (43% reduction, *p*<0.05) (Fig 6B-C). These data suggest that eNK cells can target latent reservoirs from PWH in vitro, resulting in the loss of HIV+ cells with defective proviruses as well as HIV+ cells harboring potentially intact proviruses.

### eNK cells from PWH reduce reservoirs of inducible HIV mRNA in autologous CD4 T cells after latency reversal

To quantify inducible cell-associated HIV-1 mRNA we used a next-generation sequencing-based method (EDITS assay). Latently infected cells express virtually no HIV RNA, due to the absence of HIV Tat. Once reactivated cells express a minimum level of HIV Tat, a feedback loop results in a high level of HIV mRNA accumulation. Due to this binary nature of HIV transcription, a calibration curve of the number of HIV *env* cDNA molecules detected by EDITS allows us to estimate the number of HIV *env* mRNA^+^ cells. EDITS has been used to reproducibly detect inducible *env* mRNA in latently infected T cells from well-suppressed HIV^+^ donors [65, 66]. The efficacy of a variety of LRAs has been demonstrated with this assay, including responses to HDACi, PKCa, and TCR stimulation [66]. As shown, the combination of SAHA with IL-15 or N-803 are particularly potent stimuli of HIV-1 transcription, comparable to TCR stimulation via α-CD3/α-CD28-coated beads (S5C Fig).

Using the EDITS assay to measure loss of cells expressing inducible HIV mRNA we assessed whether autologous eNK cells can reduce HIV reservoirs *in vitro* in the presence of antiretroviral drugs. After reactivation of T_m_ cells from PWH with a combination of SAHA and IL-15 overnight, followed by three days of coculture with eNK cells, significant losses of HIV *env* mRNA^+^ cells were detected in 9 out of 10 donors (Fig. 7). Due to donor variability, losses of HIV *env* mRNA^+^ cells were in the range of 32-98% but were consistently reduced overall (mean 44%) relative to reactivated T_m_ cells alone. To further optimize our cell killing assay, we conducted a time course with T_m_ cells cocultured with eNK cells at an E:T ratio of 1:1 and observed significant losses of HIV *env* mRNA^+^ cells at 3 days (56% loss; p<0.05), 5 days (67% loss; p<0.05) and 7 days (63% loss; p<0.005). The only donor that displayed an increase in HIV *env* mRNA^+^ cells at 3 days of eNK cell coculture eventually lost cells with inducible HIV *env* mRNA by 5 days. Our results confirm that eNK cells show efficient killing of T_m_ cells within 3 days, with additional killing up to at least 7 days. At all time points, uninduced cells (unstim) had virtually no detectable HIV *env* mRNA. Thus, by the EDITS (RNA) and PIPL (DNA) assays we have demonstrated for the first time that autologous eNK cells can target LRA-treated patient T_m_ cells *in vitro* to significantly reduce the HIV reservoir.

**Fig 7.**
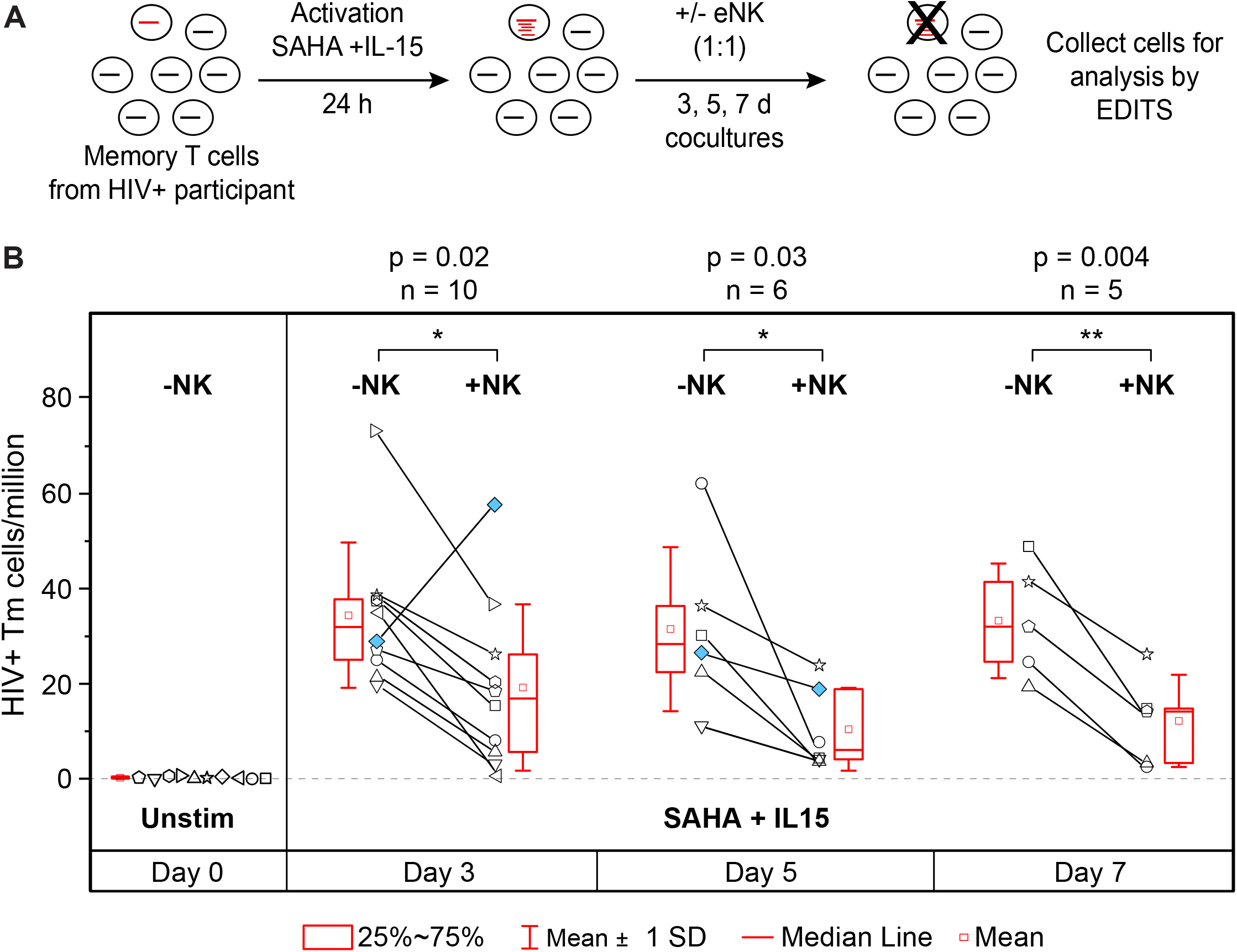
Reduction of latent HIV reservoirs in memory T cells from ART-treated PWH after latency reversal and treatment with autologous eNK cells. **(A)** Schema of the reduction of latent reservoirs in memory T cells from ART-treated PWH using eNK cells. Target cells were treated with SAHA+IL15 for 24 h, then cocultured with autologous eNK cells for 3-7 days. Killing of HIV^+^ T cells was detected as a loss of cell-associated HIV RNA by EDITS. **(B)** Box plot showing loss of HIV-infected T cells after coculture with eNK cells at 3 days (*n* = 10 biological replicates), 5 days (*n* = 6 biological replicates), and 7 days (*n* = 5 biological replicates), measured by EDITS. Each symbol represents a different donor. Statistical differences were determined using paired Student’s *t*-tests and *p* values are indicated.

## Discussion

### Expanding NK cells with mbIL-21 aAPC show cytotoxic activity for HIV-infected cells, which may provide the basis for an effective approach for an HIV Shock and Kill strategy

NK cells freshly isolated from PB are not highly cytotoxic and have limited therapeutic application. However, there have been significant advances in enhancing NK cell function for immunotherapy [67]. In this study we demonstrate that NK cells from PWH can be expanded and activated *in vitro* using aAPCs expressing membrane-bound IL-21 [48, 55, 56]. This yields a high number of CD56^bright^CD16^+^ eNK cells that express high levels of several NK cell-activating receptors, which is highly reproducible among multiple donors. Although NK cells isolated directly from peripheral blood of HIV+ donors exhibit several deficiencies relative to HIV-negative donors [45, 68–72], we have found that our expansion protocol reverses these defects. Finally, we note that an important additional advantage of using eNK cells for HIV reservoir reduction includes their high expression of Fc-γRIIIA (CD16A), which facilitates ADCC against HIV-infected cells using anti-HIV broadly neutralizing antibodies (bNAbs).

Since our goal for using eNK cells was to target acutely infected primary CD4 T cells, it was important to confirm that our highly cytotoxic eNK cells specifically target HIV-infected cells, rather than activated uninfected cells. We observed a very specific targeting of GFP^+^ HIV-infected cells with no significant death of uninfected cells. Importantly, using autologous memory T cells from PWH on ART, eNK cells significantly reduced the latent HIV reservoir after latency reversal as measured by losses in virus release, proviral load, and inducible cell-associated HIV mRNA.

There are several advantages to activating and expanding NK cells *in vitro* using aAPCs expressing membrane-bound IL-21. NK cells freshly isolated from peripheral blood can be activated temporarily *in vitro* with cytokines, which is currently in a Phase I clinical trial (NCT03346499) to assess safety after adoptive transfer. However, it is likely that a very high quantity of cytotoxic NK cells will be necessary for this approach to have any significant effect on HIV-1 reservoirs, based on previous reports for cancer treatment [73, 74]. Since the normal abundance of NK cells in peripheral blood is not sufficient for this purpose, an efficient expansion protocol will be vital. Secondly, expanded NK cells should possess a sufficiently long half-life to perform their effector function after infusion. These obstacles have been overcome by expansion of NK cells with aAPC expressing membrane bound IL-21, which generates a large number of activated NK cells that show no signs of senescence or exhaustion. aAPC-expanded NK cells are currently being developed in clinical trials for cancer therapy [48, 54–56].

### Targeting of HIV reservoirs by eNK cells requires latency reversal

There have been few studies that have shown whether a kick and kill strategy using immune effectors activated *in vitro* can significantly reduce HIV reservoirs. Primary NK cells from PWH have been activated *in vitro* with IL-15 to become cytotoxic against autologous CD4 T cells, and these activated NK cells reduced HIV reservoirs *in vitro*, as detected by quantitative viral outgrowth assay (QVOA) [75]. In a related study using LRA-treated latently infected cells, it has been shown that HIV-specific CTLs can target cells containing HIV provirus, but these effector cells failed to reduce the reservoir of replication-competent provirus that is detectable by QVOA or viral outgrowth in a murine model (MuVOA) [76]. Herzig *et al.* showed that convertible CAR-T cells can reduce the inducible latent provirus in CD4 T cells from PWH with long-term ART, after proviral reactivation with PMA and ionomycin and treatment with synthetic bNAbs fused to a MIC domain (NKG2D ligand) [77].

In our study we evaluated whether eNK cells can have a significant effect on the HIV reservoir *in vitro*, by using state of the art methods for quantifying proviruses and the potentially intact proviral load (PIPL). Traditional detection of HIV+ cells by PCR with only a single pair of primers underestimates the size of the HIV reservoir in primary PBMC samples, because of the abundance of defective proviruses that lack primer/probe binding sites. To improve sensitivity, we developed the PIPL assay using duplex digital PCR with probes for both HIV *gag* and *en*v. Since these markers are separated by ∼ 5kb in the HIV genome, we speculated that this would also allow us to detect proviruses that are potentially intact. Indeed, we discovered that detection of proviruses that are positive for both markers (*gag*+*env*+) tends to exclude defective proviruses. This is also consistent with the recent report of an intact proviral DNA assay (IPDA) using duplex digital droplet PCR [63, 64]. By counting proviruses that may have otherwise been uncounted using a single primer pair, our total proviral load counts using multiple primer pairs were inherently more sensitive.

We also evaluated the impact of eNK cells on inducible cell-associated HIV mRNA by the EDITS assay. The major advantage of using our EDITS assay to quantify the HIV reservoir is that this directly detects inducible, latently infected CD4 T cells, which produce partially spliced HIV env mRNA transcripts that are required for infectious virus production. The high sensitivity of the assay and strong enrichment for cells that may release replication-competent viruses, allowed us to demonstrate NK cell targeting of HIV reservoirs in a wide range of well suppressed donors.

If using a shock/kick and kill strategy *in vivo*, LRAs such HDACi or PKC agonists would be present in patients during adoptive transfer with effector cells. Thus, we evaluated the effect of HDACi (SAHA, panobinostat, romidepsin) and PKC agonists (bryostatin, prostratin, and ingenol) on eNK cell-mediated cytotoxicity *in vitro*. Treatment of eNK cells with bryostatin and prostratin (but not ingenol) dramatically reduced cytotoxicity, which suggests a strong inhibition of an effector function that would be essential for reducing reservoir levels in vivo. However, treatment of eNK cells with HDACi and ingenol was relatively well-tolerated. We are continuing to evaluate other LRAs for their effect on NK cell function. Notably, SAHA does not inhibit activated NK cell function when used at a physiological dose and IL-15 (or the IL-15 superagonist N-803) activates and helps maintain NK cells [67, 68]. Since we have also shown that SAHA and IL-15 synergize to promote HIV latency reversal, we propose that the combination of these agents has strong potential as a clinical regimen prior to eNK cell therapy.

Finally, successful HIV eradication will require specific targeting of the major reservoir of HIV in lymph nodes, specifically in follicular T helper cells (Tfh) in the germinal centers (GCs) of B cell follicles, and tissue-specific reservoirs in the gut and brain. The ability to genetically engineer eNK cells offers the potential to introduce homing receptors such as CCR7 and CXCR5 that can enhance entry of these cells into these immune-privileged sites.

## Materials and Methods

### Ethics Statement

The University of Pittsburgh Institutional Review Board, University Hospitals Cleveland Medical Center Institutional Review Board, and Case Western Reserve University Institutional Review Board have approved the protocols used in this study. The clinical research staff completed enrollment procedures and all participants provided written informed consent before inclusion in the study. Annually, patients in the HIV clinic at University Hospitals of Cleveland are asked if they would like to donate a blood sample to the CFAR repository. Patients who are interested are consented and PBMC and plasma samples are stored for future HIV research.

### Human subjects

For this study leukapheresis packs from 5 HIV-1-negative donors were purchased from AllCells. For HIV+ participants, PBMCs from well-suppressed ART-treated patients from the University of California, San Francisco (UCSF) SCOPE cohort and the Rustbelt Center for AIDS Research (CFAR) repository were used. The UCSF SCOPE cohort patient characteristics were a median age 40 (30-54) years, median nadir CD4 count 211 (76-661) cells/μl, median duration of viral suppression 37 (19-131) months, and duration of HIV infection range of 5-16 years [65]. The Rustbelt CFAR repository patient characteristics were a median age 50 (30-61) years, median nadir CD4 count 211.5 (0-531) cells/μl, median CD4 count 887 (492-1296) cells/μl, and a median duration on ART treatment was 8 (3-21) years [78]. For assessment of eNK cells killing of CD4+ T cells by virus release three HIV^+^ participants from the University of Pittsburgh AIDS Center for treatment were selected for leukapheresis with the following criteria: suppressive ART regimens for ≥ 24 months with CD4 counts >300 cells/µl within the past 12 months, with no active coinfections with HBV or HCV and no known malignancies.

### Molecular clones of HIV-1

The following HIV molecular clones were used: single round HIV-1 proviral clone containing *tat, rev, env, vpu*, and mCD8a-GFP-IRES-*nef* (PHR’ NL4-3-mCD8a-eGFP-IRES-Nef) [66] and full-length replication-competent HIV-1 molecular clones containing eGFP-IRES-*nef* (pNL4-3-GFP-IRES-Nef (NLgNef) and pNLAD8-GFP-Nef (AD8gNef) [79, 80].

### Cell lines

K562 ATCC CCL-243 were maintained with RPMI-1640 (Gibco Fisher Scientific) supplemented with 10% FBS (Gemini Bio-Products) and normocin (0.1 mg/ml) (InvivoGen). K562 were transduced with pRSI9-U6-(sh)-UbiC-TagRFP-2A-Puro vector (Cellecta) pseudotyped with VSV-G and selected for puromycin resistance to make K562-RFP cell line. Latent HIV-infected Jurkat E6.1 cell line (3C9) [81], which has a single insertion of HIV-1 proviral clone containing *tat, rev, env, vpu*, and d2eGFP-IRES-*nef* (PHR’ NL4-3-d2eGFP-IRES-*nef*), was maintained in RPMI-1640 supplemented with 10% FBS and normocin (0.1 mg/ml). HIV was reactivated in 3C9 cells with TNF-α(PeproTech) for 24 h. The aAPC lines C9.K562-mbIL21 and NKF were maintained in RPMI-1640 supplemented with 10% FBS and normocin (0.1 mg/ml).

### ADCC antibodies

The following reagent was obtained through the NIH HIV Reagent Program, Division of AIDS, NIAID, NIH: anti-HIV-1 gp120 monoclonal antibodies: PGT121 (ARP-12343, contributed by International AIDS Vaccine Initiative)[82], PGT126 (ARP-12344, contributed by International AIDS Vaccine Initiative)[82], PGT128 (ARP-13352, contributed by International AIDS Vaccine Initiative)[82], 10-1074 (ARP-12477, contributed by Dr. Michel Nussenzweig)[23], 2191 (ARP-11682, contributed by Dr. Susan Zolla-Pazner)[83], VRC01 (ARP-12033, contributed by Dr. John Mascola)[84], G54W (NIH45-46 G54W, ARP-12174, contributed by Dr. Pamela Bjorkman)[85], VRC03 (ARP-12032, contributed by Dr. John Mascola)[84], CH106 (ARP-12566, contributed by Drs. Barton F. Haynes and Hua-Xin Liao), 3BNC117 (ARP-12474, contributed by Dr. Michel Nussenzweig)[23], VRC-CH31 (ARP-12565, contributed by Drs. Barton F. Haynes and Hua-Xin Liao), PG9 (ARP-11557, contributed by International AIDS Vaccine Initiative)[86], PG16 (ARP-12150, contributed by International Aids Vaccine Initiative)[86], PGT145 (ARP-12703, contributed by International AIDS Vaccine Initiative)[82], HG107 (ARP-12553, contributed by Duke Human Vaccine Institute, Duke University Medical Center)[87], and CH58 (ARP-12550, contributed by Drs. Barton F. Haynes and Hua-Xin Liao); anti-HIV-1 gp41 monoclonal antibodies: 7B2 (ARP-12556, contributed by Drs. Barton F. Haynes and Hua-Xin Liao), 7B2-AAA (ARP-12557, contributed by Duke Human Vaccine Institute, Duke University Medical Center)[88], 2F5 (ARP-1475, contributed by DAIDS/NIAID)[89], 7H6 (ARP-12295, contributed by Drs. Jinghe Huang, Leo Laub, and Mark Connors), and 10E8 (ARP-12294, contributed by Dr. Mark Connors)[90]. PGT121, PGT128, PGT135 and PGT145 (IAVI NAC) were also obtained from IAVI NAC. Anti-human Influenza hemagglutinin monoclonal (CH65, ARP-12555, contributed by Drs. Barton F Haynes and Hua-Xin Liao) was used as a negative control antibody for ADCC and was obtained through the NIH HIV Reagent Program, Division of AIDS, NIAID, NIH.

### Leukapheresis processing

PBMCs were isolated from fresh leukopaks by Ficoll-Paque density gradient centrifugation. PBMCs were cryopreserved in aliquots of 50 million cells/ml, frozen using a Gordinier Controlled Rate Freezer, and stored in liquid nitrogen at −150°C until use.

### Enrichment of CD4+ T cells and memory CD4+ T cells and acute HIV-1 infection

Cryopreserved PBMCs were thawed, washed, and resuspended in RPMI 1640 medium supplemented with 10% FBS, normocin (0.1 mg/ml), and IL-2 (30 IU/ml) (PeproTech). PBMCs were incubated for two hours at 37°C prior to cell isolation. Memory CD4 T cells were isolated from PBMCs using the EasySep Memory CD4+ T cell enrichment kit (StemCell technologies). Total CD4 T cells from PBMCs were isolated using the EasySep Human CD4+ T cell enrichment kit (StemCell technologies). Prior to infection, T cells were either activated with PHA-P (5 μg/ml) (Millipore Sigma) or IL-15 (10 ng/ml) (PeproTech). T cells were infected with either replication-competent HIV-1 (pAD8gNef [80] or pNLgNef [79] or single-round HIV-1 (pHR-CD8a-d2eGFP-IRES-Nef) using Lenti-X Accelerator (TaKaRa) following the manufacturer’s protocol.

### Enrichment and expansion of NK cells

NK cells were activated and expanded as previously described with C9.K562-mbIL21 [48, 55] or NKF cells [56]. Briefly, NK cells were either isolated from cryopreserved PBMCs by either negative selection using the EasySep NK cell enrichment kit (StemCell technologies) or by depletion of T cells with human CD3 MicroBeads (Miltenyi Biotec). NK cells were cultured in SCGM (CellGenix) supplemented with 10% FBS, normocin (0.1 mg/ml) and IL-2. aAPC were γ-irradiated at 100 Gy and then cryopreserved until ready for use. NK cells were expanded with either C9.K562-mbIL21 (at a 2:1 ratio (aAPC:NK cells) for the first stimulation and 1:1 ratio for subsequent stimulations in the presence of 100 IU/ml of IL-2 or with NKF (at a 5:1 ratio (aAPC:NK cells) for the first stimulation and a 2:1 ratio for subsequent stimulations in the presence of 120 IU/ml of IL-2. Irradiated cells were added to expanding NK cells cultures once every 7 days for approximately 3-4 weeks.

### Assessment of NK cell function

#### Surface marker expression

For surface staining of NK cells, about 1-2 x 10^5^ cells were stained with fluorescent antibodies for 20-30 min at room temperature, washed, and resuspended in PBS. Data was acquired using a BD LSR Fortessa (BD Biosciences) and analyzed using WinList 3D 9.0.1 (Verity Software House). The following anti-human antibodies used were obtained from BD Biosciences: CD56 (NCAM16.2), CD3 (HIT3a), NKp30 (p30-15), and NKp44 (p44-8), CD16 (3G8), NKG2D (1D11), NKp46 (9E2), DNAM1 (DX11), 2B4 (2-69), and CD57 (NK-1), from R&D Systems: NKG2A (131411), NKG2C (134591), KIR3DL1 (177407), and KIR2DL1 (143211); and from Miltenyi Biotec KIR2DL2/DL3 (DX27).

#### Flow cytometry NK killing assays

Killing of target cells by NK cells was done using different assays. For all the killing assays, data was acquired using a BD LSR Fortessa and analyzed using WinList 3D 9.0.1.

The PanToxiLux (OncoImmunin, Inc.) assay measures killing of target cells by NK cells and was set up following the manufacturer’s protocol. Briefly, target cells (K562 or CD4 T cells) were stained with TFL4 to discriminate between T cells and NK cells and NFL1 to differentiate dead cells prior to coculture with NK cells. In some experiments, target cells were not stained with TFL4 but instead NK cells were stained with 1 μM CellTrace Violet (Cell Proliferation dye, ThermoFisher Scientific) following the manufacturer’s protocol to help differentiate between target cells and effector cells. Cocultures of target cells and eNK cells were established with different eNK cell inputs and incubated for 1 h in the presence of a fluorogenic granzyme B and caspase 8 substrate (PS). The PanToxiLux assay measures target cells that have received a lethal dose of granzyme B from NK cells after coculture, which is detected by PS cleavage.

NK cell cytotoxicity was also measured by their ability to kill K562-RFP target cells. K562-RFP cells were cocultured with different inputs of CellTrace Violet-stained NK cells. To determine the effect of LRAs on NK cell cytotoxicity, NK cells were treated with LRAs (330 nM SAHA (Cayman Chemical), 10 nM romidepsin (Selleck Chemicals), 20 nM panobinostat (Selleck Chemicals), 5 nM bryostatin (Sigma-Aldrich), 1 µM prostratin (Sigma-Aldrich) and 100 nM ingenol (Santa Cruz Biotechnology)) for 24 h prior to staining with CellTrace Violet. Killing of K562-RFP cells leads to loss of RFP and it directly correlates with gain of a dead cell stain (Fixable Viability Dye, eBioscience). Dead cells were stained following the manufacturer’s protocol.

Killing of HIV-1-infected CD4 T cells was also detected by loss of GFP and gain of a dead cell stain (propidium iodide, PI (Cayman Chemicals). To achieve an optimal percentage of (HIV)GFP+ cells, acutely infected T cells were mixed with autologous uninfected T cells, followed by dead cell removal (Dead Cell Removal Kit, Miltenyi Biotec). NK cells were stained with CellTrace Violet as indicated above. For killing of CD4 T cells acutely infected with replication-competent HIV, cocultures of CD4 T cells and NK cells were established at 1:1 and 1:10 (NK:target) inputs in a 96-well plate and incubated for 24 h in SCGM supplemented with 10% FBS, normocin, and IL-2 (100 IU/ml). After 24 h coculture, dead cells were stained with 5 µg/ml PI for 15 min. Cells were then washed and fixed with 4% formaldehyde (methanol-free, Polysciences). In this assay, killing of HIV(GFP+)-infected CD4 T cells was measured by loss of GFP and killing of all cells was measured by gain of PI stain. Percent killing is shown as number of GFP+ CTV-PI-from cultures with NK cells divided by GFP+ CTV-PI-from cultures without NK cells. All killing assays were carried out in triplicate for each sample.

#### NK cell killing detected by microscopy

Dead cells were removed from target cell cultures and CellTrace Violet-stained NK cells were added at a 1:10 (NK:T cell) ratio. Sytox Orange Dead Cell Stain (1 µM) (ThermoFisher Scientific) was immediately added to CD4 T and NK cell culture and cells were loaded into a hemacytometer. Cells were visualized using a DeltaVision Core Deconvolution microscope, equipped with an LED light source and an environmental chamber to maintain conditions of 37° C and humidified CO2 that are optimal for cell viability. Time lapse images were acquired at 30 second intervals for 45 minutes per sample using a CCD camera, 40X lens, and SoftWoRX Pro 6.5.1 software(Applied Precision).

### eNK cell killing assay against autologous CD4+ T cells from PWH for HIV detection by EDITS, proviral load measurement and virus release

Leukapheresis from ART-treated HIV+ participants were collected and processed as described above. NK cells were enriched from PBMCs by CD3 depletion (Miltenyi) and expanded in presence of irradiated NKF cells as described above. CD4 T cells were isolated from autologous donors and treated with LRA for 24hr. When HIV was reactivated using Dynabeads Human T activator CD3/CD28 for T cell expansion and activation (ThermoFisher Scientific), beads were magnetically removed the next day. When HIV was reactivated with SAHA (500 nM) and IL-15 (10 ng/ml) or N-803 (1 nM) (Altor Bioscience) combination, LRAs were left in the culture. For killing assays that required virus release measurements, large cocultures of 20-40 million unstimulated or stimulated (LRA-treated) T cells with an equal number of eNK cells (1:1 ratio of NK:total CD4 T cells) were established in upright T75 flasks. For killing assays were only the EDITS and proviral load was the required for HIV detection, cocultures of 2-3 million CD4 T cells and equal number of eNK cells (1:1 ratio of NK:total CD4 T cells) were established. All cocultures were maintained in SCGM supplemented with IL-2 (120 IU/ml) and in the presence of 10 µM raltegravir (Sigma-Aldrich) and 0.4 µM nevirapine (Sigma-Aldrich) for 3, 5, or 7 days. For large cocultures were supernatant was collected for virus release detection, one third to half the media from the top of the culture was collected, centrifuged at 600 x g for 10 min, supernatant was then transferred to another tube and flash frozen in a dry ice/ethanol bath. Frozen supernatants were kept at −80 °C until ready for HIV release analysis. Fresh media supplemented with IL-2 and antiretrovirals were added back to cocultures. At 5 day coculture, a complete media change was performed and cells were resuspended back in fresh media supplemented with IL-2 and antiretrovirals. After 7 day coculture, NK cells were removed from cultures (EasySep Human CD56 Positive Selection Kit II, StemCell Technologies). Remaining CD4 T cells were analyzed by flow cytometry to confirm removal of NK cells by staining for CD56, CD3, and CD4. For the EDITS and Proviral assays, 1-2 million viable remaining CD4 T cells were flash frozen in a dry ice/ethanol bath and stored in a liquid nitrogen tank until ready for analysis.

#### Detection of inducible cell-associated HIV-1 mRNA by EDITS

We performed the EDITS assay as previously described [65] with the following modifications. Briefly, memory CD4 T cells were activated for 24 h with the LRA(s) indicated. When specified, eNK cells added to T cell cultures were removed (EasySep Human CD56 Positive Selection Kit II, STEMCELL Technologies) prior to analysis. A total of 1 million viable T cells were pelleted for each sample to be analyzed at the time points indicated, followed by removal of supernatant and freezing of cell pellets in a dry ice and ethanol bath. For RNA isolation, cells were lysed at 55° C for 10 min in 75 µl of lysis buffer [0.1 M Tris pH 8.0, 0.2 M NaCl, 5 mM EDTA, 1% SDS, and 0.2 mg/ml Proteinase K (ThermoScientific)] with shaking at 1500 rpm. After the lysis, 25 µl of 100% isopropanol and 180 µl of Cytiva Sera-Mag SpeedBeads Carboxylate-Modified Magnetic Particles (Fisher Scientific) (suspended in 2.5 M NaCl, 1 mM trisodium citrate, 20% PEG 8000, 0.05% Tween-20, pH 6.4) were incubated with RNA for 5 minutes and immobilized with a magnet. RNA-bound beads were washed twice with Washing Buffer I (0.5% SDS, 80% EtOH, 1 mM trisodium citrate, pH 6.4) and once with Washing Buffer II (80% EtOH, 1 mM trisodium citrate, pH 6.4). RNA-bound beads were then DNase treated (10 mM Tris-HCl [pH 8], 2.5 mM MgCl2, 0.5 mM CaCl2, and 1 Unit RNase-free DNase) for 15 min at 37° C. A solution of 20% PEG 8000 and 2.5 mM NaCl was added to the DNase-treated RNA-bound beads. Beads were then washed twice with Washing Buffer II and dried. RNA was eluted in 40 µl of 10 mM Tris pH 8.0. One quarter of the total RNA sample was used as template for OneStep RT-PCR (ABM) using the following primers: EDITS Fwd 5’-GTTGTGTGACTCTGGTAACTAG-3’ and EDITS Rvs 5’ CTGAAGATCTCGGACCCATTGT-3’ and cycling conditions: 42° C for 30 min, 94° C for 3 min, 35 cycles of (94° C for 30 s, 55° C for 30 s, 72° C for 60 s), and 72° C for 5 min. A nested PCR was then performed using 4% of OneStep RT-PCR product mixed with BestTaq 2x Master Mix and barcoded primers containing linkers compatible with Ion Torrent Sequencing and the following HIV-1-specific sequences: EDITS nFwd 5’-TAGTCAGTGTGGAAAATCTCTA-3’ and EDITS nRvs 5’-CATAATAGACTGTGACCCACAA-3’. Nested PCRs were amplified with the following conditions: 94° C for 3 min, 5 cycles of (94° C for 10 s, 60° C for 30 sec, 72° C for 10 sec), 30 cycles of (94° C for 10 s, 72° C for 15 s), and 72° C for 3 min. Nested PCR samples were then pooled, purified by agarose gel electrophoresis, quantified with Qubit HS DNA Assay and analyzed by next generation sequencing using the Ion Chef System and Ion S5 System (ThermoFisher Scientific) following the manufacturer’s protocol. Sequences were then mapped to an HIV genome reference and quantified using Geneious Prime software (Geneious).

#### HIV Potentially Intact Proviral Load (PIPL) assay by duplex digital PCR

Cellular DNA from memory or total CD4 T cells of HIV-positive donors was isolated using a Qiagen DNeasy Blood & Tissue kit, eluted in Qiagen EB Buffer, and quantified using a Qubit™ dsDNA HS assay kit. Proviral loads were measured by the QuantStudio™ 3D Digital PCR System (Life Technologies). 300-700 ng of cellular DNA (per chip) was mixed with QuantStudio™ 3D Digital PCR Master Mix v2 and the following primer/probe sets in duplex: 300 nM *gag* forward primer 5’-AGCCCAGAAGTAATACCCATGTTT*-*3’ (HXB2 position 1282-1305), 300 nM *gag* reverse primer 5’-CCCCCCACTGTGTTTAGCATG*-*3’ (HXB2 position 1347-1367), 300 nM *env* forward primer 5’-TGGAAAAATGACATGGTAGAACAGA-3’ (HXB2 position 6510-6534), 300 nM env reverse primer 5’-TTTACACATGGCTTTAGGCTTTGA-3’ (HXB2 position 6563-6586), 300 nM TaqMan® *gag* probe 5’-(6-FAM)-CAGCATTATCAGAAGGAGCC-(MGBNFQ)-3’ (HXB2 position 1307-1326), and 300 nM TaqMan® *env* probe *5’*(VIC)-CATGAGGATATAATCAGTTTATGG-(MGBNFQ)-3’ (HXB2 position 6537-6560). PCR mixes were loaded in 2-3 replicates onto QuantStudio® 3D Digital PCR 20K Chips v2 using a QuantStudio® 3D Digital PCR Chip Loader.

Sealed chips were cycled using a Dual Flat Block GeneAmp™ PCR System 9700 (Applied Biosystems®) at 96°C for 10 minutes, followed by 40 cycles of 60°C for 2 min and 98°C for 30 sec, with a final extension at 60°C for 2 min. Fluorescence was measured with a QuantStudio 3D Digital PCR Instrument and analyzed using Analysis Suite dPCR Cloud Software (Thermo Fisher Connect™ Platform). HIV-specific calls in FAM (*gag*) and VIC (*env*) channels were assigned relative to background from an HIV negative donor (No-Amp). Call numbers were converted to copy numbers based on a modified multiplicity of infection equation, P(n) = e^-m^m^n^/n!, where n = copy # per well, and m = copy # / available well #. Available well # = loading efficiency x 20,000 wells/chip x # chips. PCR volume analyzed (based on loading efficiency) equals available well # x 0.755 nL/well. Wells containing a detectable copy of target sequence are “calls” but underestimate copy number due to Poisson distribution, because any well with >1 copy creates an undercount. The probability that each well with PCR mix contains 2 to 7 detectable copies [P(n)] was used to estimate this undercount, which was added back to call # to yield copy #. Data from multiple chips are combined during analysis to improve confidence. The number of human cells represented in each PCR mix was based on the concentration of cellular DNA and mass of the human diploid genome (6.6×10^-12^*g* DNA/cell).

#### Measurement of HIV-RNA (virus release)

Supernatants from eNK-treatment cultures were tested for HIV-RNA using an Aptima HIV-1 Quan Dx assay kit on the Hologic Panther® platform (Hologic). When multiple test values for a single sample could not be averaged due to non-detectable or unquantifiable results, those which were non-detectable were assigned a value of ½ the lower limit of detection (LLoD=12cp/mL) and those which were detectable but unquantifiable were assigned a value of ½ the lower limit of quantitation (LLoQ=30cp/mL).

### Statistics

Statistical analysis was determined by a Student’s paired *t* test using Origin (OriginLab Corporation) or GraphPad Prism. *P* values less than 0.05 were considered significant. Figure legends indicate the number of independent experiments or individual donor samples.

## Acknowledgements

We would like to acknowledge Dr. Dean Lee for contributing the C9.K562.mbIL21 feeder cells used in our NK cell expansion protocols. We thank Jennifer Bongorno Hurt for NLgNef and AD8gNef virus preparation. The BD LSR Fortessa and the DeltaVision Deconvolution microscope Core facilities were supported by the National Institutes of Health Grant P30 AI036219 to Case Western Reserve University/University Hospitals Center for AIDS Research. The irradiation of cells was supported by the Radiation Resources Core Facility of the Comprehensive Cancer Center of Case Western Reserve University and University Hospitals of Cleveland (P30 CA43703). We thank study participants for the donation of leukapheresis units.

## Author Contributions

M.C. designed and implemented the experiments, performed data analysis, and prepared the manuscript. B.L. designed PIPL DNA assay and K562-RFP cell killing assay, performed time-lapse microscopy and cytotoxicity experiments (PIPL/K562), and contributed to data analysis and manuscript preparation. C.D. performed the EDITS assays. D.N.W. provided NKF cells. D.M. and G.H. developed clinical protocol, regulatory oversight, and participant enrollment. M.D.S. and P.N.E. performed the measurement and analysis of virus release and J.C. contributed to virus release data analysis. J.K. and J.W.M. contributed to the concept of the work, the design of experiments, data analysis and critical review of the manuscript.

## Supporting Information Captions

**S1 Fig.**
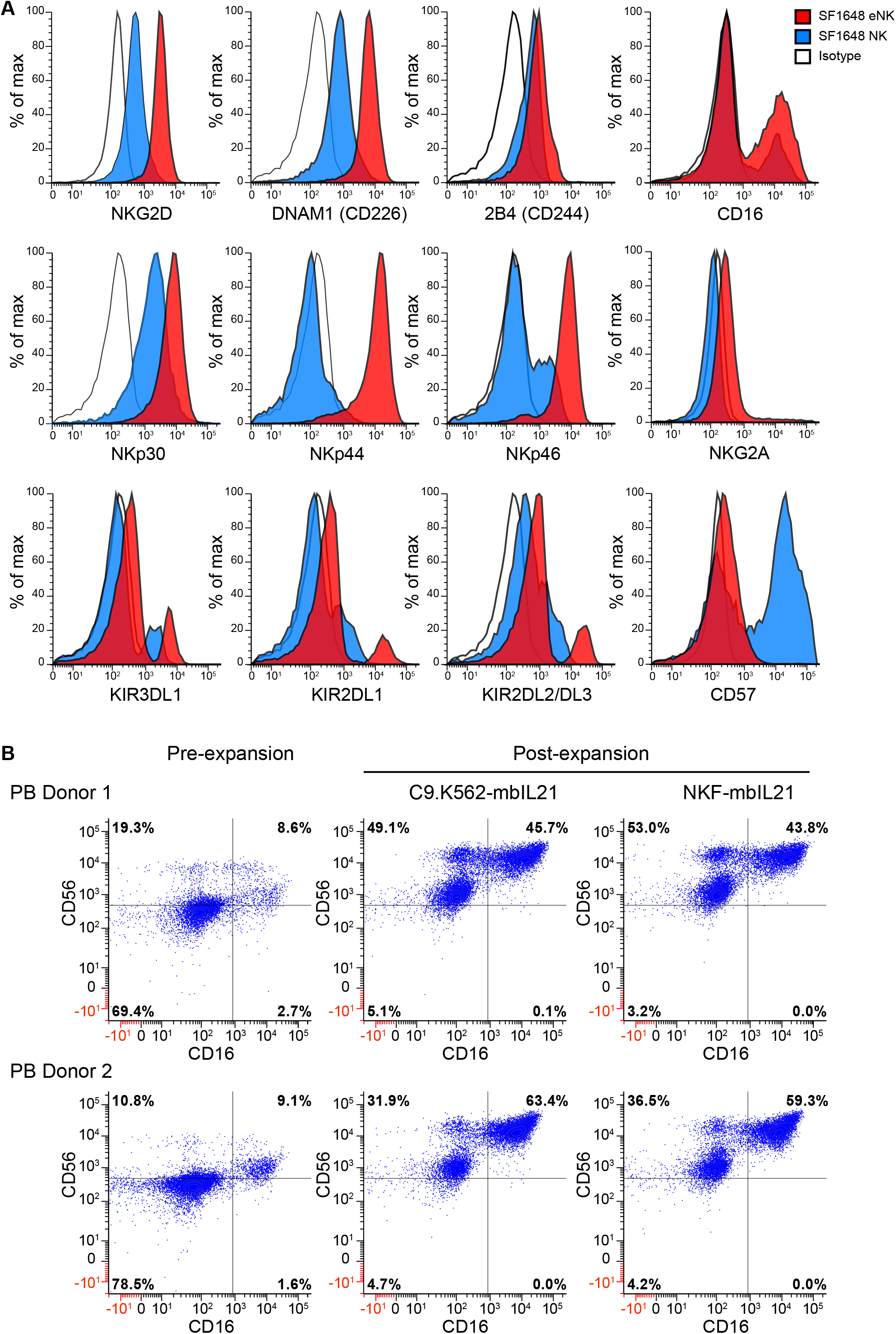
Expansion of NK cells from HIV+ participants yields CD56^bright^ eNK cells that express high levels of activating receptors. **(A)** Flow cytometry analysis showing MFI for various NK cell receptors of NK cells from a representative HIV+ participant before expansion (SF1648 NK in blue) and after expansion (SF1648 eNK in red). Isotype is shown in white. **(B)** eNK cell phenotype analysis by flow cytometry using two different artificial antigen presenting cells expressing membrane-bound IL-21. Dot plots gated on CD3-cells showing expression of NK cell markers CD56 and CD16 before and after (pre- and post-) expansion using irradiated K562.C9-mbIL21 cells or irradiated NKF-mbIL21 cells. (n = 2 biological replicates)

**S2 Fig.**
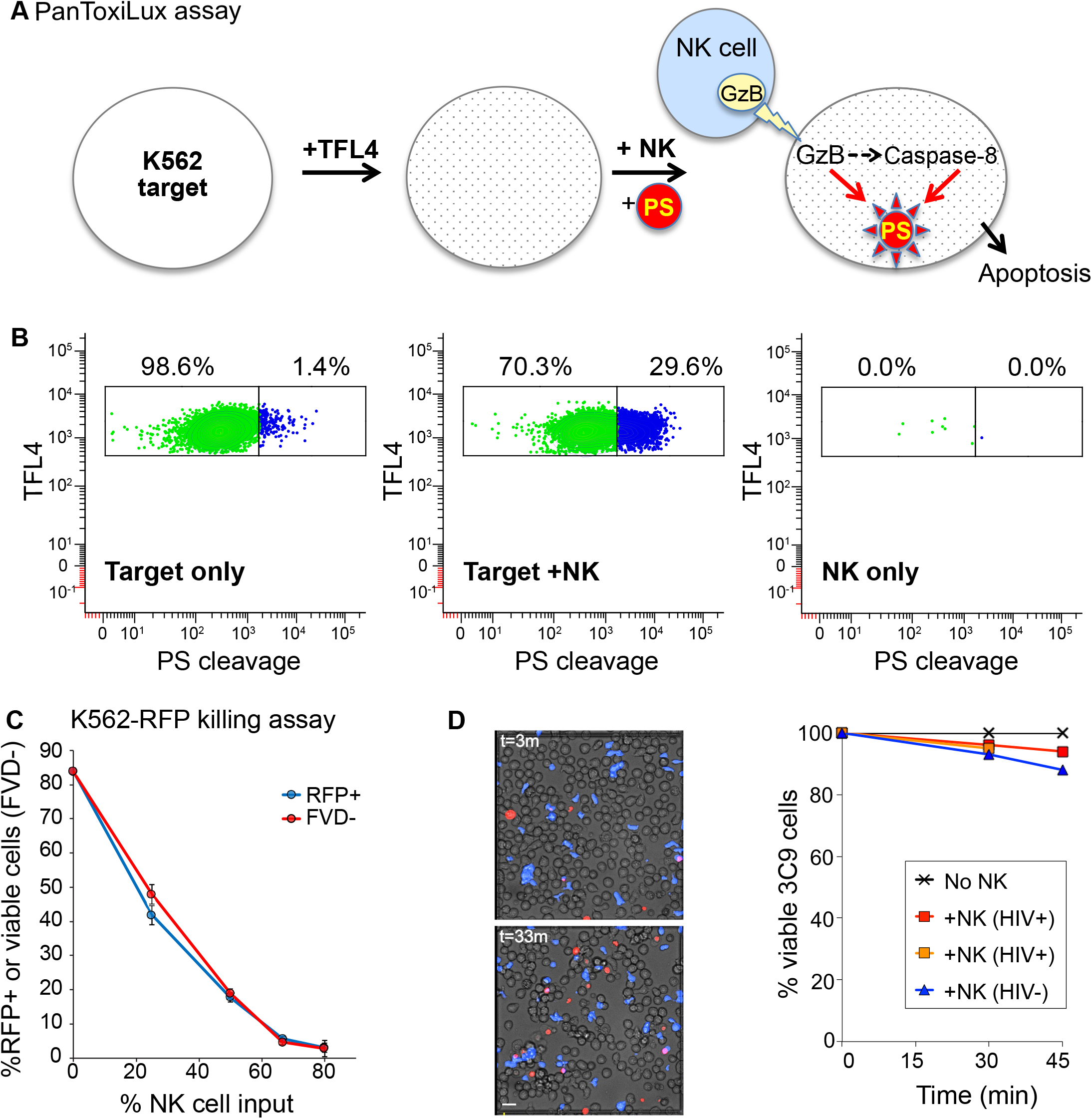
Flow cytometry assays to detect eNK cell-mediated killing of K562 target cells. **(A)** Schematic representation depicting cell cytotoxicity assay (PanToxiLux, OncoImmunin). This assay utilizes a unique cell-permeable granzyme B/Caspase-8 substrate (PanToxiLux Substrate, PS), which becomes fluorescent upon cleavage in target cells. NK cells kill target cells by delivering granzyme B (GzB) into the cytoplasm, which can also activate Caspase-8, resulting in fluorescence when PS is present. Target cells are stained with a fluorescent dye (TFL4) to differentiate them from NK cells and with a fluorescent dead cell dye (NFL1) to eliminate target cells that have died prior to the assay. Both TFL4 and NFL1 are intracellular stains that do not interfere with NK cell recognition. Stained target cells are washed to remove excess dye and cocultured with NK cells for 1 h in the presence of PS. **(B)** Flow cytometry gating strategy for the PanToxiLux cytotoxicity assay. Dot plots are gated on TFL4+NFL1-events to exclude NK cells and dead cells. TFL4+ target cells with PS cleavage (blue) vs. lack of PS cleavage (green) are shown. Dead K562 cells are shown as being TFL4+ NFL1-PS cleavage+. **(C)** Flow cytometry NK cell cytotoxicity assay using K562 constitutively expressing RFP (K562-RFP cell line). CFSE-stained NK cells and K562-RFP target cells are cocultured overnight at different NK:K562 ratios. By gating for CFSE-negative K562 cells, cytotoxicity is detected by loss of RFP and quantified as a ratio of (initial/final) %RFP cells. Dead cells were stained with a fixable viability dye (FVD). The percentage of live cells (FVD-) is directly proportional to RFP+ cells. **(D)** Time lapse imaging of eNK cells from an HIV+ donor cocultured with TNF-α-treated 3C9 cells (HIV+ Jurkat cells). TNF-α-treated 3C9 cells were mixed with a dead cell stain and with fluorescently CTV-labeled eNK cells (blue) from either HIV- or HIV+ donors, then added to a hemocytometer and imaged by time-lapse microscopy for 45 min. eNK and target cells were cocultured at a 1:10 (9% NK input) (NK:target cell) ratio. Nuclei of dying cells are stained in real time with Sytox Orange Dead Cell Stain (red). Representative images at 3 and 33 min are shown (left). Quantitation of dead 3C9 cells after coculture with eNK cells (right). Viable and dead 3C9 cells were counted at 0, 30, and 45 minutes. Percent viable 3C9 cells at 0 minutes are normalized to 100% and the number of dead cells over the total of viable cells at 0 minutes are shown in the graph.

**S3 Fig.**
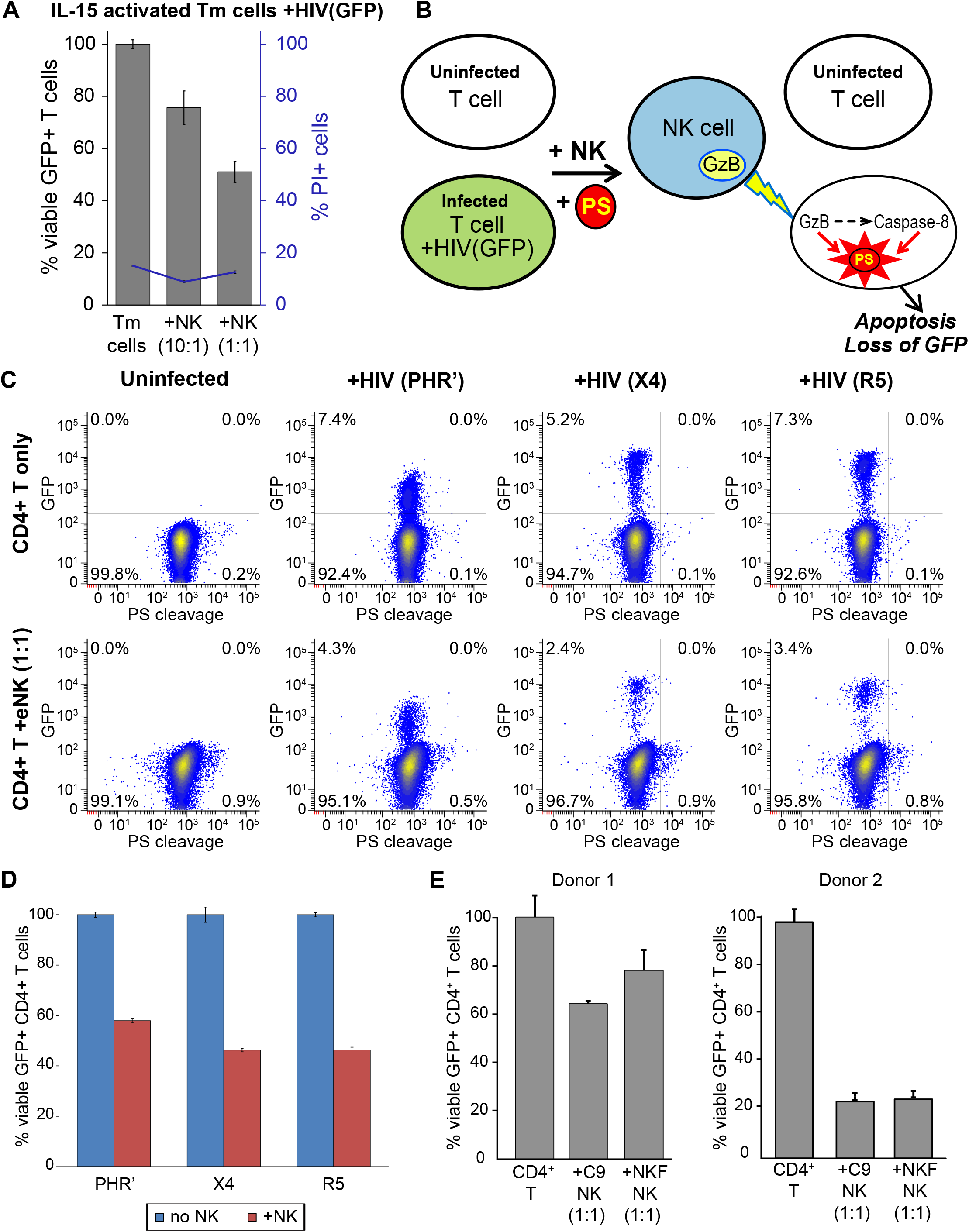
Specific eNK cell killing of primary T cells infected with HIV. **(A)** Quantitation of viable HIV-infected (GFP+) T_m_ cells remaining after coculture with autologous eNK cells. IL15-activated memory T cells (T_m_) were acutely infected with R5-tropic HIV(GFP). CTV-labeled NK cells were cultured with autologous HIV(GFP)-infected Tm for 1 day, and dead cells were then stained with PI and analyzed by flow cytometry. The percentage of viable GFP^+^ T_m_ cells (gated on CTV^-^ PI^-^ cells) cultured with eNK cells was normalized to cell cultures without eNK cells. The percentage of total dead CD4 T cells (CTV-%PI+ cells) is shown in blue. **(B-D)** eNK cell-mediated killing of HIV-infected CD4 T cells using an adapted PanToxiLux assay. **(B)** Schematic of adapted PanToxiLux assay to detect killing of HIV(GFP)-infected cells. Specific killing of HIV(GFP)+ cells leads to loss of GFP, due to loss of plasma membrane integrity, while killing of any target cell (HIV+/-) is measured by PS cleavage. NK cells are fluorescently labeled with CTV to differentiate them from T cells. **(C)** Activated CD4+ T cells were infected with replication-competent X4 or R5-tropic HIV(GFP) or transduced with a VSV-G pseudotyped HIV construct (PHR’) that contains *tat*, *rev*, *env*, *vpu*, *nef* and mCD8α-GFP. Uninfected and infected target cells were cocultured with autologous eNK cells (CTV+ stained) at a 1:1 ratio. Representative dot plots are shown and are gated on CTV-events for killing analysis. Viable HIV-infected T cells are GFP+ PS cleavage-. Viable uninfected T cells are GFP-PS cleavage-. **(D)** Analysis of killing of HIV-infected CD4 T cells by autologous eNK cells. Bar graph shows the relative percentage of GFP+ cells that are viable after coculture with eNK cells. The percentage of viable GFP+ T cells is normalized to viable GFP+ T cells in cultures without eNK cells. *n* = 1 biological replicate with each sample performed in triplicate. **(E)** eNK cell cytotoxicity against HIV-infected T cells after expansion with two different aAPC expressing membrane-bound IL-21. eNK cell killing of autologous primary CD4+ T cells acutely infected with R5-tropic replication-competent HIV-GFP. NK cells expanded with C9.K562.mbIL21 (C9-NK) or NKF.mbIL21 (NKF-NK) were CTV-stained and cocultured at a 1:1 eNK:T cell ratio. Killing of HIV-infected T cells is measured by loss of GFP in CTV-cells by flow cytometry. Bar graph shows the percentage of viable GFP+ cells after coculture with autologous eNK cells, normalized to viable GFP+ T cells cultured alone (*n* = 3 replicates per donor). Error bars represent SEM.

**S4 Fig.**
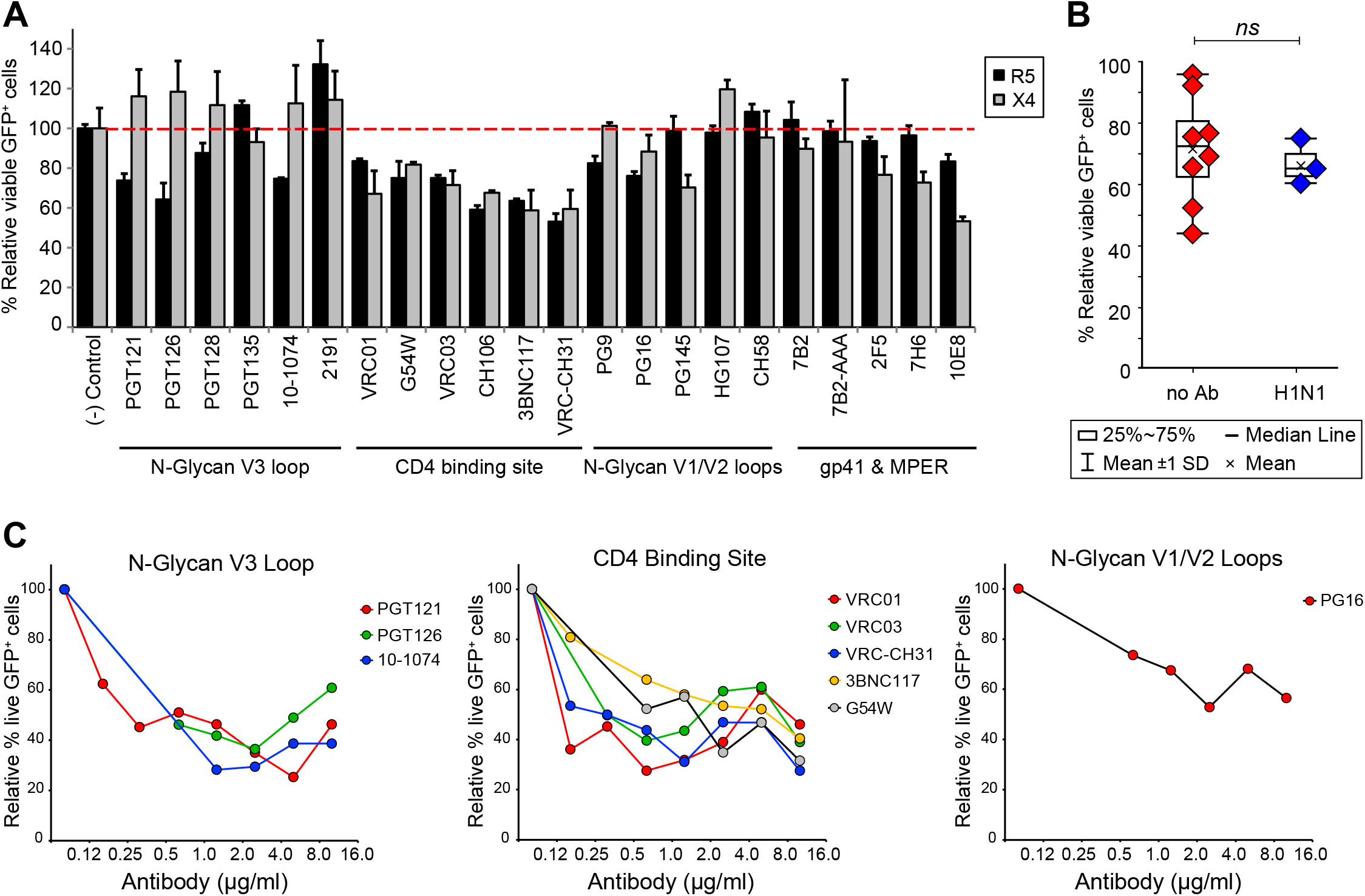
Enhanced specificity of eNK cells against HIV-infected cells by ADCC using bNAbs. **(A-C)** eNK cells were pre-stained with CTV and cocultured with autologous, HIV(GFP)-infected CD4 T cells at an NK:T cell ratio of 1:10 in the presence of individual antibodies (10μg/ml) and raltegravir for 1 day. Cells were then stained with PI and analyzed by flow cytometry. **(A)** Screening of anti-Env bNAbs for ADCC using eNK cells and R5 or X4-tropic HIV(GFP)-infected CD4+ T cells. Bar graph shows the relative percentage of GFP+ cells that are viable after coculture with eNK cells. The percentage of viable GFP+ T cells is normalized to viable GFP+ T cells in cultures with a negative control, which was either human anti-influenza A virus (H1N1) hemagglutinin for R5 or with no antibody for X4. bNAbs that elicited ADCC show killing below the red dashed line. (*n* = 1 biological donor in triplicate) **(B)** Anti-H1N1 antibody does not elicit ADCC by eNK cells against HIV-infected CD4 T cells. For no Ab *n* = 4 biological replicates with each symbol representing the result from an independent experiment performed in triplicate, and for anti-H1N1 *n* = 1 biological replicate with each symbol representing the result from an independent experiment performed in triplicate **(C)** Titration of ADCC candidate bNAbs. eNK and HIV-infected CD4 T cells were cocultured as described above in the presence of bNAbs at 0.125-10 µg/ml and analyzed by flow cytometry. Error bars represent SEM.

**S5 Fig.**
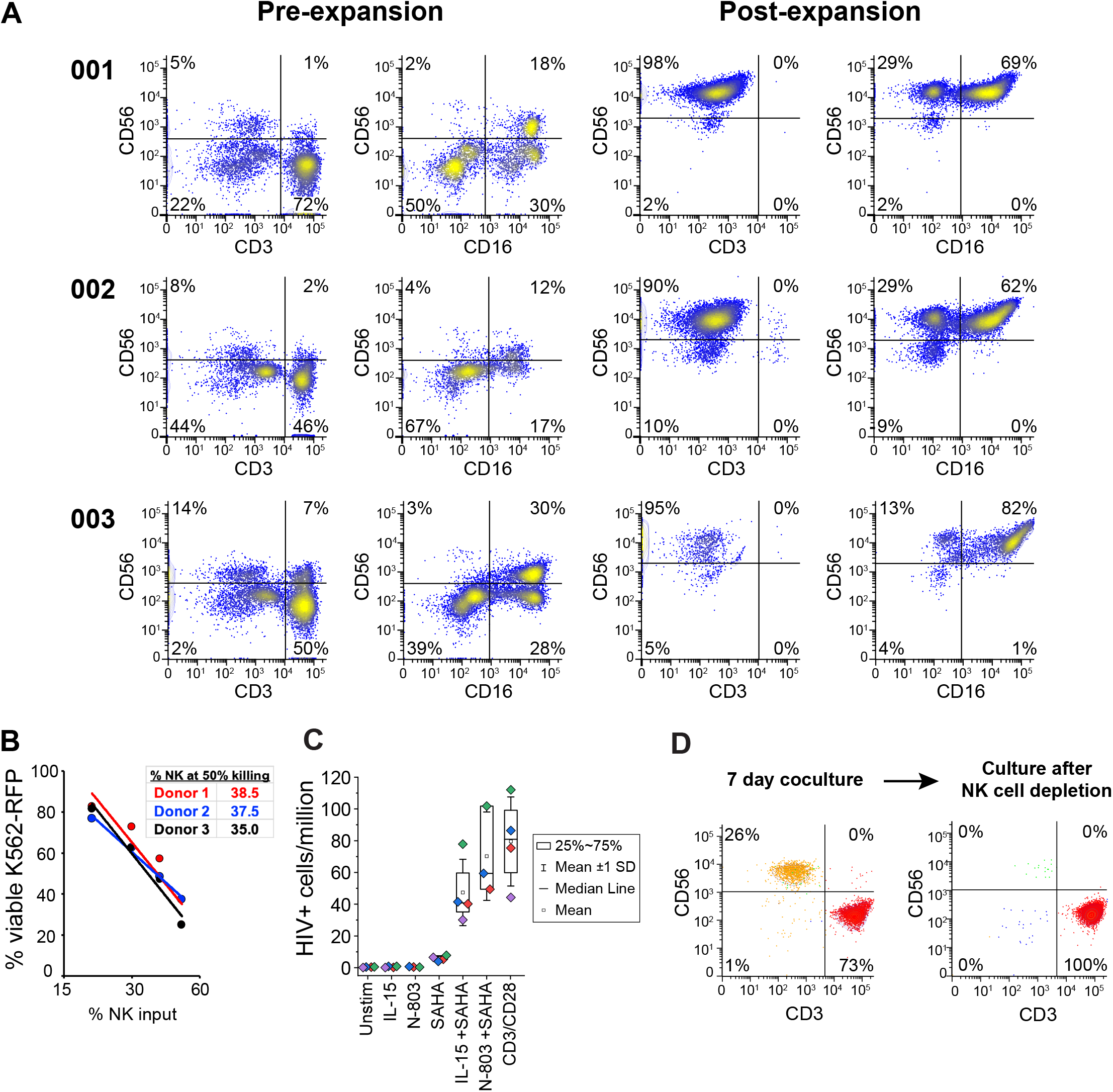
Reduction of virus release after LRA treatment and coculture with GMP-like eNK cells. **(A)** NK cell phenotypic analysis by flow cytometry before and after expansion using irradiated GMP-like NKF cells (aAPC expressing mbIL21) for Donors 001-003. Dot plots showing the proportion of NK cells and T cells before and after (pre- and post-) expansion, as well as the expression of NK cell markers CD56 and CD16 (gated on CD3-) for each donor are shown. **(B)** Flow cytometric GMP-eNK cell cytotoxicity assay using K562-RFP cell line for Donors 001-003. CFSE-stained NK cells and K562-RFP target cells were cocultured overnight at different NK:K562 ratios. By gating on CFSE-negative events, cytotoxicity is detected by loss of % viable K562-RFP. For comparison among donors, % NK cell input at 50% killing is also shown. (n = 3 biological replicates) **(C)** EDITS assay to detect HIV mRNA+ cells in CD4 T cells from ART-treated PWH. CD4 T cells were activated for 24 h with various LRAs indicated on the box plot. The number of HIV-1 mRNA+ cells per million CD4 T cells is shown (n = 4 biological replicates). **(D)** Removal of NK cells by CD56 positive selection. Staining for CD56 and CD4 of eNK and CD4 T cells in a representative coculture at Day 7 before and after NK cell removal (culture after NK cell depletion).

**S6 Fig.**
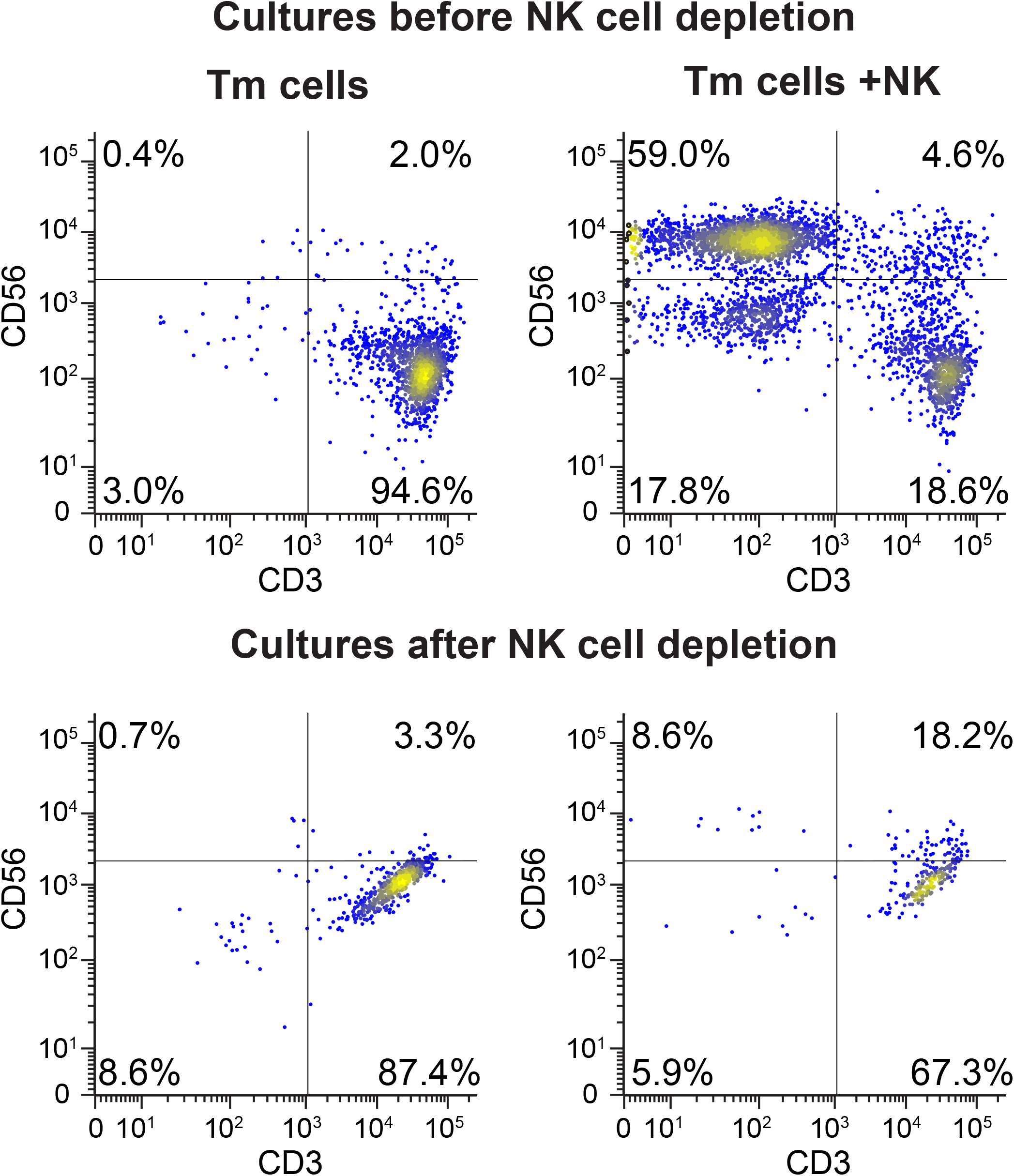
Efficiency of NK cell depletion after coculture of eNK cells with CD4 T cells. After 7 days culture of SAHA+IL15-treated memory T cells alone (T_m_ cells) or with NK cells (+NK), cells were stained with anti-CD56 (NK cell marker) and anti-CD3 (T cell receptor) before (top) and after (bottom) CD56+ NK cell depletion. After NK cell depletion, DNA from T_m_ cells was isolated for proviral load analysis.

### Multimedia Files

**S1 Movie. Time lapse imaging of TNF-α-treated 3C9 cells (Negative control).** 3C9 cells were treated with TNF-α for 24 h and then added to a hemacytometer slide in presence of Sytox Orange Dead Cell Stain (red) for real time staining of nuclei in dying cells. The time-lapse imaging is performed at 37 °C and begins immediately after addition of cells to the slide and lasts for 45 min.

**S2 Movie. Time lapse imaging of eNK cells from an HIV+ donor cocultured with TNF-α-treated 3C9 cells.** 3C9 cells were treated with TNF-α for 24 h. eNK cells were pre-stained with CTV proliferation dye (blue). eNK and 3C9 cells were mixed at a 1:10 (9% NK input) (NK:target cell) ratio in presence of Sytox Orange Dead Cell Stain (red) and then added to a hemacytometer slide. The time-lapse imaging is performed at 37 °C and begins immediately after addition of cells to the slide and lasts for 45 min.

